# A neuron-glia circuit anticipates hypoxia to regulate organismal oxygen use

**DOI:** 10.64898/2026.04.10.717666

**Authors:** Rongwei Zhang, Ziqiang Wei, Javier J. How, Michele Nardin, Sujatha Narayan, Amina Kinkhabwala, Weiyu Chen, Jing-Xuan Lim, Virginie M. S. Ruetten, Anuththara Rupashinge, Martin Haesemeyer, Brett D. Mensh, Mark C. Fishman, Florian Engert, Behtash Babadi, Jiulin Du, David A. Prober, Misha B. Ahrens

## Abstract

Organisms must regulate metabolic resources such as oxygen (O_2_) and nutrients despite environmental variability and the energetic costs of their own actions^1–3^. Such regulation can occur reactively, through homeostatic corrections of recent imbalances, or predictively, through allostatic adjustments that anticipate future demand^4,5^. Predictive regulation is particularly important because metabolic resources often continue to be consumed for seconds to minutes after motor actions cease as tissues repay incurred costs, making it advantageous to prevent depletion before it occurs^6^. However, the cellular and circuit mechanisms for allostatic control remain largely unknown^5,7,8^. Using whole-brain neuronal and astroglial imaging and O_2_ measurements in behaving zebrafish, we identified a noradrenergic–astroglial circuit that detects, anticipates, and prevents internal O_2_ depletion. We found that swimming exacerbated internal hypoxia with a multi-second delay, but behavioral adaptations occurred before such self-generated hypoxia manifested, suggesting predictive control, confirmed using computational modeling. Noradrenergic neurons in the nucleus of the solitary tract directly detected brain hypoxia and received efference copies of swimming actions; these inputs summed at the level of membrane voltage to increase spiking and norepinephrine release when actions and resource scarcity co-occurred. Astroglia integrated noradrenergic input into prolonged Ca^2+^ elevation that tracked the O_2_ cost of recent actions and thereby predicted O_2_ debt relative to O_2_ availability, rising ~8 s before O_2_ fell. This astroglial prediction reorganized brain-wide activity to suppress locomotion and promote respiration, preempting O_2_ depletion. Silencing noradrenergic neurons or astroglial signaling abolished these hypoxia coping behaviors, whereas selective activation evoked them. This neuronal–astroglial mechanism constitutes a predictive control system that integrates physiological state with behavioral intent to avert metabolic crisis, revealing a cellular substrate for proactive energy management.

## Main

Animals must regulate their physiology and behavior in response to changing environmental and internal resources^9–14^. Both aquatic and terrestrial animals consume nutrients and O_2_ as they run, swim, or fly^15–18^, while also experiencing varying resource availability caused by environmental^19–21^ and bodily factors^22^. Survival under stressors like hypoxic conditions, such as fish in stagnant water^23,24^, thus requires brain mechanisms to estimate the physiological risk of performing vigorous physical actions when resources are limited, and regulate bodily O_2_ levels for future use (Fig. 1a). Motor actions rapidly consume ATP, but consume O_2_ gradually with a multi-second lag^6,25^, requiring that effective behavioral regulation be based on both current reserves and anticipated future O_2_ depletion. Thus, although animals might in principle just react to current internal O_2_ levels, an alternative and more effective strategy would be to also make a prediction of how internal O_2_ levels will vary in the future as a result of recent actions and preemptively adjust behavior to minimize anticipated hypoxic stress^3,8,26^. However, little is known about behavioral control strategies that predict the physiological risk of actions and their cellular and circuit-level mechanisms.

**Figure 1.**
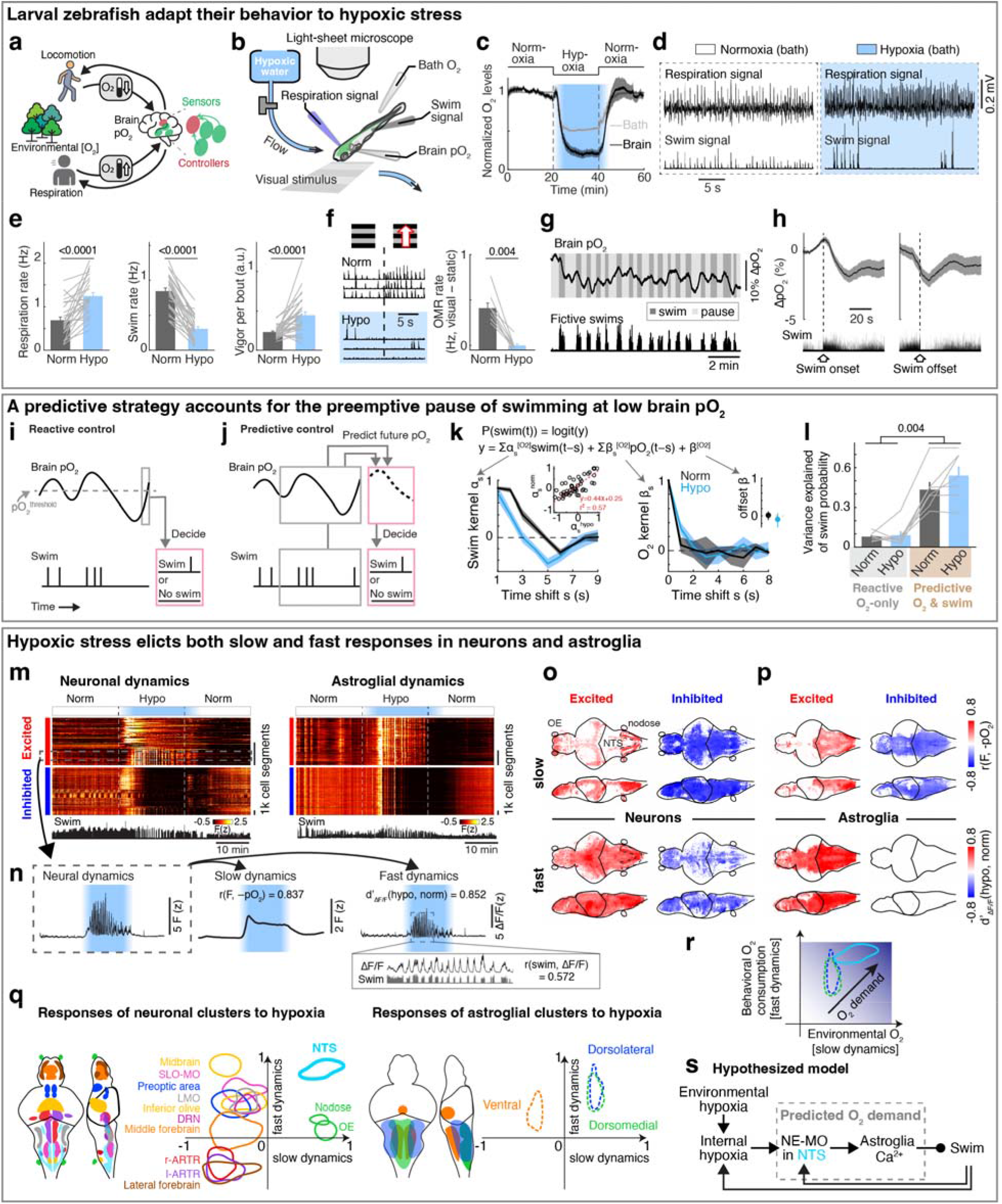
Larval zebrafish show oxygen-dependent changes in behavior and brain-wide calcium dynamics. **a**, Hypothetical schematic showing the interactive relationship between brain O_2_ levels (pO_2_) and behavior. **b**, Experimental schematic showing measurements of fictive respiration and fictive swimming using glass electrodes placed on the head and tail, brain and bath O_2_ levels, optional visual stimulus, and brain-wide calcium imaging using a light-sheet microscope. **c**, Mean ± s.e.m. brain (black) and bath O_2_ (gray) levels during normoxia and hypoxia for 8 fish. The O_2_ values were normalized to the mean during normoxia. **d**, Electrophysiological recording of fictive respiration and swimming in a paralyzed fish during normoxia and hypoxia. **e**, Mean ± s.e.m. fictive respiration and swimming during normoxia and hypoxia for 29 paralyzed fish (gray lines). Hypoxia leads to more respiration and less frequent, stronger swims. Wilcoxon signed-rank test. **f**, Optomotor response (OMR) to forward visual flow during normoxia and hypoxia for 9 paralyzed fish. OMR rate is calculated by subtracting swim rate during visual stimulus to that during static period. Wilcoxon signed-rank test. **g**, Spontaneous fluctuations of brain pO_2_ and fictive swimming in a paralyzed fish. See Extended Data Fig. 1i for non-paralyzed fish. **h**, Normalized brain pO_2_ aligned by swim onset or offset from paralyzed fish (134 swim events from 12 fish). Gray shadows indicate s.e.m. **i**, Schematic of a reactive model where a swim action depends on current brain pO_2_. Dashed gray line, brain pO_2_ threshold for swimming; pink boxes, transform model from current brain pO_2_ (top gray box) to the probability of a swim action (bottom pink box). **j**, Schematic of a predictive model where a swim decision depends on (1) recent brain pO_2_ over a prolonged period (top gray box), and (2) recent swim actions (bottom gray box) from which self-generated hypoxia is estimated; pink boxes, transform model from the internal estimate of future brain pO_2_ (dashed line, top pink box) to the probability of a swim action (bottom pink box). **k**, Model fits of the predictive model to behavioral data during normoxia (black) and hypoxia (blue). Top, a generalized linear model for the predictive model. Bottom left, fitted swim kernel; bottom right, fitted oxygen kernel; right inset, offset; left inset, linear correlation between α_s_ in hypoxia and normoxia conditions; black circles, α_s_ from the same fish; red line, linear fit. Mean ± s.e.m. is shown for 7 fish. **l**, Variances explained by the reactive (pO_2_-only) and predictive (pO_2_ & swim) models. Bar graphs indicate mean ± s.e.m. across 7 fish; two-way ANOVA based on the model type and O_2_ condition. Variances explained by the reactive model were 0.08 ± 0.03 (mean ± s.d., normoxia) and 0.09 ± 0.07 (hypoxia). Those by the predictive model were 0.43 ± 0.14 (normoxia) and 0.54 ± 0.17 (hypoxia). **m**, Example activity matrices showing oxygen-responsive neurons (left) and astroglia (right) from one *Tg(elavl3:H2B-jGCaMP7f)* fish and one *Tg(gfap:GCaMP6f)* fish, which are detected using rank correlation between the inverse brain pO_2_ (−pO_2_) and the calcium dynamics of individual cells. Red bar: hypoxia-excited cells. Blue bar: hypoxia-inhibited cells. The cell segments are sorted based on unsupervised factor-based clustering. More clusters and average spatial distributions for additional fish are shown in Extended Data Fig. 2c-f. Bottom black traces show fictive swimming; vertical scale bars, 1k (1000) cell segments. **n**, Raw fluorescence of a neuronal cluster indicated by the boxed region in (m) was decomposed into slow and fast dynamics. Their relationships to hypoxia were quantified as r(F, −pO_2_), the rank correlation with −pO_2_, and d’_ΔF/F_(hypo, norm), the d-prime comparison of ΔF/F between the final 10-minutes of normoxia and hypoxia. r(swim, ΔF/F) measures rank correlation between the instantaneous swim vigor and ΔF/F. Panels indicate values of these measures for the example neuronal cluster. **o**,**p**, Whole-brain maps showing hypoxia-excited and -inhibited slow (top) and fast (bottom) components for neurons in 9 fish (o) and astroglia in 6 fish (p). **q**, Brain schematics showing the location of 10 neuronal (left) and 3 astroglial (right) clusters (see Extended Data Fig. 2c,e for supporting data) and joint distribution plots of each cluster’s slow (x-axis) and fast (y-axis) dynamics to hypoxia. Positive and negative values indicate hypoxia-excited and -inhibited cells, respectively. The circle-like shapes represent the kernel density estimate of 70% of cells in each cluster from 9 fish (neurons) and 6 fish (astroglia). Notably, cells in the NTS cluster (cyan) uniquely show hypoxic excitation in both the slow and fast components. **r**, Hypothetical schematic showing that oxygen demand depends on environmental O_2_ levels and behavior-related O_2_ consumption; the increase of blue gradient indicates an elevation of total O_2_ demand. The solid cyan, dashed blue, and dashed green regions represent the calcium responses of NTS neurons, dorsolateral astroglia, and dorsomedial astroglia, respectively, to hypoxia, as shown in (q). **s**, Hypothetical circuit schematic for hypoxia-induced effects on brain cell dynamics and behavior. Arrow, excitation; line ending in filled circle, inhibition. Blue shading indicates hypoxic conditions in (c,d,e,f,k,l,m,n) and all following figures.

### Zebrafish larvae adapt behavior to varying oxygen levels

To examine the effects of hypoxia on the brain and behavior, we established a preparation in larval zebrafish that included whole-brain calcium imaging of neurons and astroglia, while monitoring fictive swimming using electrophysiology, and measuring O_2_ levels in both the water bath and in the brain using Clark-type O_2_ microelectrodes with sub-second response times^27,28^ (Fig. 1b; see Methods). In addition, using separate recording electrodes placed on the skin above the head, we also captured respiration-relevant electrical signals, which were synchronized with bodily movements associated with respiration^29^, including movements of the operculum, jaw, and pectoral fins (Extended Data Fig. 1a-c). Perfusion of a hypoxic solution (4 mg/l O_2_ compared to 8 mg/l O_2_ for normoxia) resulted in a drop of O_2_ levels by ~50% in the bath and ~80% inside the brain by ~5 minutes (Fig. 1c), a difference that is likely due to consumption of O_2_ by the animal.

Similar to hypoxia-induced adaptive behaviors that have been described in many species^30,31^, acute hypoxia resulted in an increase of respiration rate, a decrease of swim frequency with an increased occurrence of long pauses (5 seconds or more of no swimming^32^), more vigorous swim bouts, and decreased behavioral responsiveness to visual stimuli (Fig. 1d-f and Extended Data Fig. 1d-h). These effects can be viewed as an adaptive response wherein the fish minimize the decrease in brain O_2_ level (pO_2_, partial pressure of oxygen) during hypoxia by decreasing energy expenditure (less swimming) while increasing O_2_ intake (more breathing).

### Reactive versus predictive control reflected by fish behaviors

In addition to the effects of environmental O_2_ levels, swimming caused a ~2% per 10 seconds decrease in brain pO_2_ (Fig. 1g and Extended Data Fig. 1i,j). This decrease lagged multiple seconds behind locomotion, continuing for several seconds after swimming ceased, and recovered on a timescale of ~10 seconds (Fig. 1g,h and Extended Data Fig. 1i-k). The effect was observed both during fictive swimming in paralyzed fish and in restrained, non-paralyzed fish (Extended Data Fig. 1i), consistent with O_2_ consumption arising from neuronal activity in the brain and spinal cord^33–35^. The decrease was larger in non-paralyzed fish, presumably due to muscle contractions.

To mitigate the effects of hypoxia, one possible behavioral strategy is to stop swimming when brain pO_2_ drops below a threshold, a form of reactive control (Fig. 1i). However, because pO_2_ continues to decline for several seconds after swimming ends—as it is consumed by oxidative phosphorylation to regenerate metabolic resources like ATP in muscle and nerve cells^6,36,37^, and as O_2_ equilibrates diffusively throughout tissues^38^ (Fig. 1h and Extended Data Fig. 1j)—fish might also benefit from proactively adjusting behavior in anticipation of such delayed O_2_ depletion. This would use an alternative predictive control strategy, in which recent swimming history is used to estimate forthcoming O_2_ decline and preemptively pause locomotion (Fig. 1j and Extended Data Fig. 1l,m). These two strategies—reactive and predictive—provide complementary hypotheses for how behavioral control of O_2_ is organized.

To test which strategy best describes fish behavior, we fitted swim data with two models (Fig. 1i,j): (1) a reactive controller in which swim actions depend only on current pO_2_, and (2) a predictive controller in which fish integrate both recent pO_2_ and past swim actions to estimate future pO_2_ and select subsequent actions. The predictive model outperformed the reactive model with a high level of variance explained (>40%), and explained fourfold more variance in behavior (Fig. 1k,l; see Methods). The fitted kernels revealed two key components: (1) a direct dependence on current pO_2_ (Fig. 1k, right; reactive component), and (2) an hypoxia-dependent integration of swims (Fig. 1k, left; predictive component). The swim-integration weights during hypoxia summed to a larger value than those during normoxia. Moreover, the swim kernel was biphasic: swims initially promoted more swimming but suppressed it after ~3 seconds, generating a “struggle then pause” pattern. Under hypoxia, the net effect was inhibitory, agreeing with preemptive pauses.

Together, these analyses defined the computational requirements of a predictive controller: a representation of current brain pO_2_, an integrator of recent motor output that predicts future pO_2_ decreases, a multiplicative interaction between these signals to represent expenditure relative to availability, and downstream control of swimming and respiration to preemptively compensate for O_2_ debt before it manifests. Guided by this theoretical framework, we next examined the neural substrates implementing predictive O_2_ regulation.

### Neuronal and astroglial representations of brain pO_**2**_

In order to identify the control mechanism through which prediction of pO_2_ regulated behavior, we performed whole-brain light-sheet calcium imaging of neurons and astroglia in response to changes in pO_2_ (Fig. 1m; *Tg(elavl3:H2B-jGCaMP7f)* fish for neurons and *Tg(gfap:GCaMP6f)* fish for astroglia). Based on the predictive model, we hypothesized that adaptations to hypoxia are driven by brain activity that accounts for two factors on different timescales: a slow factor on the timescale of tens of seconds reflecting changes in pO_2_ that tend to arise from ambient environmental O_2_ level fluctuations, and a fast factor on the timescale of seconds reflecting O_2_-consuming swim behaviors. We therefore decomposed brain-wide neuronal and astroglial calcium signals into a slow component (F_slow_) which mainly reflected pO_2_-related baseline calcium dynamics, and a fast component (ΔF/F) which captured behavior-related calcium transients (Fig. 1n and Extended Data Fig. 2a; r(swim) represents rank correlation between swim vigor and ΔF/F; see Methods).

Next, we related F_slow_ to brain hypoxia (−pO_2_) using rank correlation (Extended Data Fig. 2b), and compared fast dynamics (ΔF/F) during hypoxia and normoxia using d-prime (see Methods). Acute hypoxia treatment induced dynamic responses in both neurons and astroglia (Fig. 1m and Extended Data Fig. 2c-f; identified via factor-based clustering, see Methods). For slow and fast dynamics, we considered cells to be ‘hypoxia-excited’ if their calcium activity increased during the hypoxic period, i.e. showing positive correlation to −pO_2_ for F_slow_ and increased d-prime during hypoxia for ΔF/F, respectively (red in Fig. 1o,p), or ‘hypoxia-inhibited’ if calcium activity decreased during hypoxia (blue in Fig. 1o,p). Additionally, hierarchical clustering of calcium activity identified several populations that showed more complex slow dynamics (e.g., a biphasic response to hypoxia; Extended Data Fig. 2b). For slow dynamics, hypoxia-excited neurons were distributed sparsely in the brain, primarily located in the olfactory bulb, anterior forebrain, optic tectum, lateral hypothalamus, nucleus of the solitary tract (NTS), and nodose ganglia (Fig. 1o, top, red). In contrast, hypoxia-inhibited neurons were observed in most brain areas (Fig. 1o, top, blue), likely due to reduced ATP generation and thus reduced energy available for spiking during hypoxia^39^. Hypoxia-excited astroglia were widespread but most prominent in the dorsal hindbrain, including the NTS and surrounding region, whereas hypoxia-inhibited astroglia were distributed more widely in the forebrain, hypothalamus and anterior hindbrain (Fig. 1p, top). Distinct from hypoxia-induced modulation of neuronal F_slow_, different neuronal clusters showed more diverse types of fast dynamics during hypoxia (Fig. 1o, bottom, and Extended Data Fig. 2c,d), whereas almost all astroglial cells showed elevated fast dynamics during hypoxia (Fig. 1p, bottom, and Extended Data Fig. 2e,f). Furthermore, the fast dynamics in many neuronal populations and all astroglial cells were synchronized with swims during hypoxia (Fig. 1m,n, and Extended Data Fig. 2c-f), suggesting that anticipated hypoxic stress caused by the combination of low pO_2_ and motor activity may be represented by specific neuronal and all astroglial populations. Interestingly, when the sodium channel blocker tricaine was added to the water bath to inhibit neuronal activity^40^, astroglial responses to hypoxia were largely suppressed (Extended Data Fig. 2g,h), which suggests that astroglial responses to hypoxia are mainly driven by neuronal activation. Overall, hypoxia induced a global reconfiguration of neural and astroglial dynamics.

Based on this analysis, neurons in the NTS and astroglia in the dorsal hindbrain stood out, as they showed the largest increases of both the slow and fast calcium components in response to hypoxia (Fig. 1q). As described above, slow dynamics can represent environmentally-induced O_2_ level changes in the brain, while fast dynamics may be related to anticipated O_2_ consumption due to behavior. These signals can be integrated as an internal estimate of future O_2_ demand (Fig. 1r), consistent with the fish following a predictive strategy (Fig. 1j), which is related to forward models in motor control^41^. In the regime of current as well as anticipated high future O_2_ demand, the brain would suppress nonessential movements to preserve O_2_. We found the majority of hypoxia-activated NTS neurons co-localized with NE-MO (the noradrenergic cluster of the medulla oblongata, likely homologous to A2 in mammals)^42^ (Extended Data Fig. 2i,j). Considering that noradrenergic neurons can activate astroglia^32,43^, we thus considered the noradrenergic-astroglial pathway as a candidate to detect brain hypoxia and modulate the behaviors that facilitate O_2_ replenishment (Fig. 1s and Extended Data Fig. 1n,o).

### Brainstem noradrenergic neurons directly detect intracranial pO_**2**_

To examine whether and how hypoxia activates noradrenergic populations in the NTS (i.e. NE-MO), we imaged *Tg(th1:Gal4);Tg(UAS:GCaMP6f)* fish, which express GCaMP6f in all noradrenergic neurons in the brainstem including NE-MO, the locus coeruleus (LC) and area postrema (AP), and sympathetic ganglia (SG, in the peripheral nervous system) (Fig. 2a). Most of these noradrenergic neurons showed hypoxia-related signals, but to different extents (Fig. 2b-d). NE-MO and SG neurons showed large F_slow_ increases during hypoxia, LC showed moderate increases, and AP showed no change (Fig. 2c,d). Moreover, hypoxia enhanced the swim-related fast dynamics (ΔF/F) in NE-MO and LC, but SG and AP neurons showed little or no change (Fig. 2c,e). Conversely, treatment with hyperoxic water (100% increase in bath O_2_ relative to normoxia) led to a decrease of F_slow_ in NE-MO, SG and LC, but not in AP (Fig. 2f). Strikingly, we found that F_slow_ in these areas correlated negatively with brain pO_2_, with strongest correlations in NE-MO and SG, less in LC, and none in AP (Fig. 2g,h). These results imply a link between brain pO_2_ and the activity of noradrenergic neurons in the brain, particularly NE-MO.

**Figure 2.**
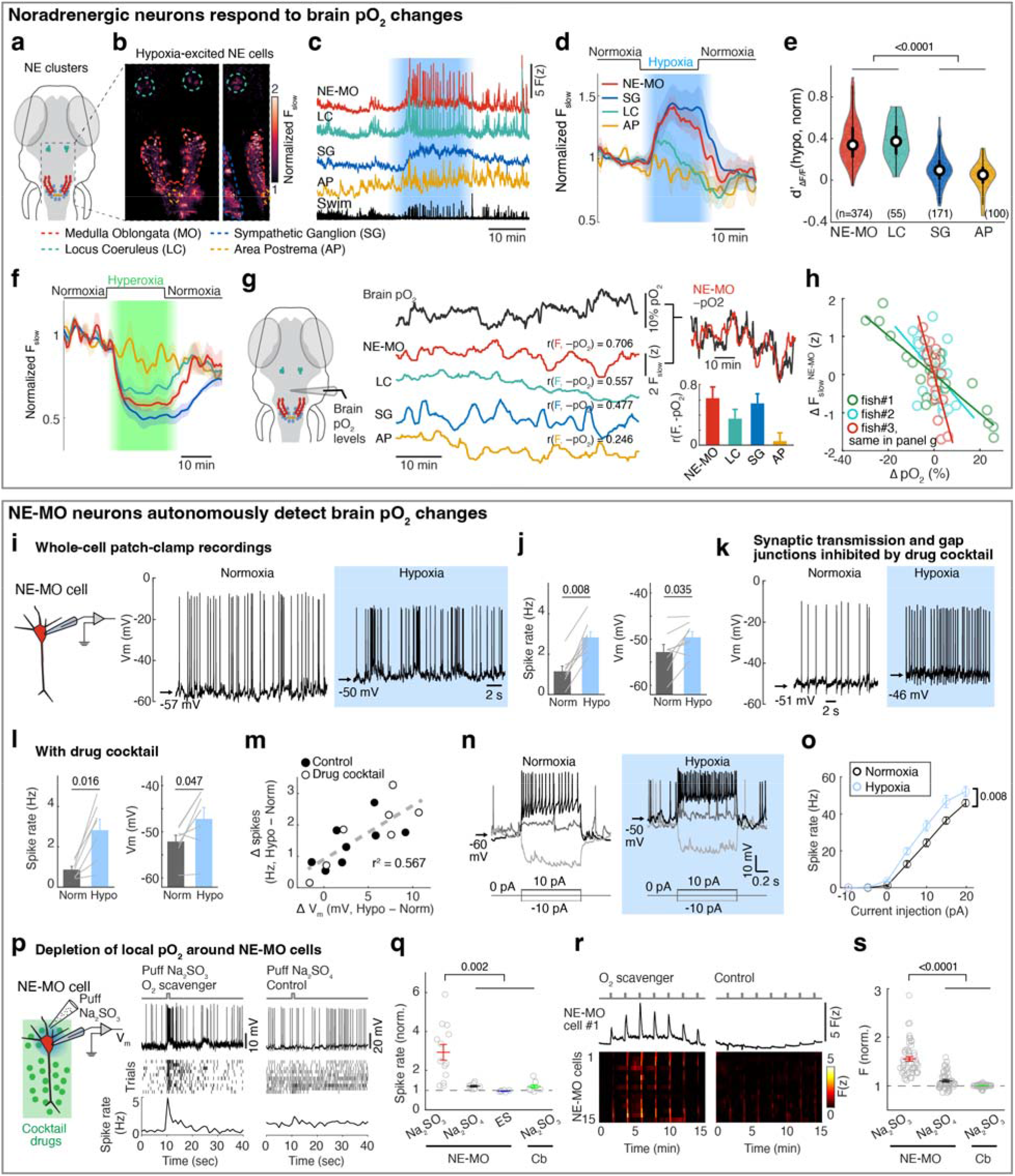
Noradrenergic neurons are directly responsive to brain pO_2_ changes. **a**, Schematic diagram showing the locations of the four groups of noradrenergic neurons. The boxed region is shown in (b). **b**, Hypoxia-excited noradrenergic neurons in *Tg(th1:Gal4);Tg(UAS:GCaMP6f)* fish. Dorsal (left) and side (right) views are shown. Normalized F_slow_ was calculated by dividing the average of slow calcium fluorescence (F_slow_) in hypoxia by that in normoxia. **c**, Example GCaMP6f fluorescent traces in the four noradrenergic populations and fictive swimming during normoxia and hypoxia. **d**, Normalized slow calcium component of the four noradrenergic neuron groups in response to hypoxia across 7 fish. Shading indicates s.e.m. **e**, Violin plots of ΔF/F amplitude change of the four noradrenergic neuron groups in response to hypoxia versus normoxia. Wilcoxon rank-sum test. The numbers in the brackets indicate the cell number from 7 fish. **f**, Normalized slow calcium component of the four noradrenergic neuron groups in the brain in response to hyperoxia across 5 fish. Shading indicates s.e.m. **g**, Simultaneous measurements of brain pO_2_ and F_slow_ of NE-MO, LC, SG and AP from an unparalyzed fish. The numbers above each trace indicate the rank correlation between brain pO_2_ and F_slow_. Right top: overlapping traces showing inverse relationship between NE-MO F_slow_ and brain pO_2_ (plotted as **−**pO_2_). Right bottom: cross-correlation analysis between brain pO_2_ and F_slow_ of noradrenergic neuron groups across 3 fish. **h**, The relationship between the changes of brain pO_2_ and NE-MO F_slow_ from 3 fish. Each dot represents the mean value over 2 minutes. Fish #3 (red) corresponds to the fish shown in panel (g). **i**, Whole-cell patch-clamp recording of NE-MO neurons and example traces showing that membrane potential and spike firing rate increase under hypoxia (see also Extended Data Fig. 4a,b). The arrows indicate the resting membrane potentials. **j**, Effects of hypoxia on resting membrane potential and spiking from 8 NE-MO neurons. Wilcoxon signed-rank test. **k**, Example traces showing the increase of membrane potential and spiking of a NE-MO cell under hypoxia while synaptic transmission and gap junctions were inhibited. **l**, Effect of hypoxia on resting membrane potential and spiking from 7 NE-MO neurons when synaptic transmission and gap junctions were inhibited. Wilcoxon signed-rank test. **m**, Scatter plot showing a positive correlation between hypoxia-induced changes in membrane potential and spike frequency in 15 NE-MO cells, 7 of which (open circles) were recorded during inhibition of synaptic transmission and gap junctions. Dashed line, a linear fit. **n**,**o**, Example traces (n) and summary (o) showing the effect of current injection on membrane potential of 8 NE-MO cells during normoxia and hypoxia. Light and dark gray lines at the bottom of (n) indicate different current injection amounts. Two-way ANOVA test. **p**, Left, schematic of local application of the O_2_ scavenger sodium sulfite (Na_2_SO_3_) or negative control sodium sulfate (Na_2_SO_4_) during whole-cell patch-clamp recording of a NE-MO cell. Right, examples showing that NE-MO cell spike firing was increased by puffing Na_2_SO_3_ but not Na_2_SO_4_. **q**, Summary of spike rate changes of NE-MO and cerebellar (Cb) neurons evoked by puff application of Na_2_SO_3_ (13 NE-MO neurons; 6 Cb neurons), Na_2_SO_4_ (8 NE-MO neurons), or extracellular solution (ES) (6 NE-MO neurons). Puff-triggered spike rates were normalized to baseline firing rates before puffing, yielding fold changes during drug application. The dashed line at one indicates the normalized baseline. Wilcoxon rank-sum test. More control experiments for puffing are shown in Extended Data Fig. 4k,l; the effects on V_m_ are shown in Extended Data Fig. 4m,n. **r**, Example traces and matrices showing the effect of puffing Na_2_SO_3_ or Na_2_SO_4_ on calcium activities of the same NE-MO cells using *Tg(th1:Gal4);Tg(UAS:GCaMP6f)* fish. **s**, Summary of calcium activity of NE-MO and Cb neurons evoked by puffing Na_2_SO_3_ (50 NE-MO neurons and 53 Cb neurons) or Na_2_SO_4_ (50 NE-MO neurons) in *Tg(th1:Gal4);Tg(UAS:GCaMP6f)* fish. The dashed line at one in indicates the normalized baseline. Wilcoxon rank-sum test.

Previous studies have shown that brain cells can sense hypoxia from specific peripheral organs^44,45^. We tested this possibility by ablating known peripheral neurons related to sensing O_2_, including the olfactory epithelia and nodose ganglia. However, neuronal, astroglial and behavioral responses to hypoxia were intact after each ablation (Extended Data Fig. 3). We therefore tested whether hypoxia can directly affect the physiology of NE-MO neurons using whole-cell patch-clamp electrophysiology (Fig. 2i and Extended Data Fig. 4a,b). First, we found that hypoxia led to an increase of resting membrane potential (V_m_) and both tonic and phasic spike firing of NE-MO neurons (Fig. 2i,j and Extended Data Fig. 4a-f). In contrast, hypoxia had only a moderate effect on LC neurons, and little or no effect on AP neurons (Extended Data Fig. 4d-f), consistent with the calcium responses of these cells during hypoxia (Fig. 2b-g). Thus, NE-MO neurons can detect brain hypoxia via an increase in both V_m_ and spike rate, and may function to modulate behavior to compensate for low O_2_ levels.

Next, we performed two experiments to determine whether NE-MO neurons can directly detect hypoxia. First, to exclude the possibility of synaptic inputs from upstream O_2_ chemoreceptors, we used a drug cocktail to abolish glutamatergic and cholinergic synaptic transmission as well as gap junction function (Methods)^46^. Treatment with this drug cocktail abolished fictive locomotion, and almost completely silenced brain-wide neuronal activity (Extended Data Fig. 4g,h) and phasic spike firing of NE-MO neurons (Fig. 2k), indicating a near-complete block of neuron-neuron communication. However, hypoxia still caused an increase in V_m_ in NE-MO neurons to the same level as without the drug cocktail (Fig. 2k-m and Extended Data Fig. 4i,j), suggesting that the response of NE-MO neurons to reduced brain pO_2_ does not depend on inputs from peripheral sensors. Rather, NE-MO cells may detect brain pO_2_ themselves and respond to hypoxia by increasing V_m_ and tonic spike firing rate, likely through intrinsic cellular mechanisms^47,48^ or through local cell-cell interactions^49,50^. Beyond the increase of V_m_ and tonic firing rate during hypoxia, we found that NE-MO cells become more responsive to input, as injecting current with the same amplitude caused more spike firing during hypoxia than during normoxia (Fig. 2n,o).

Second, we manipulated local pO_2_ by puffing the O_2_ scavenger Na_2_SO_3_ directly onto the soma of NE-MO neurons, and measured the responses of NE-MO neurons using patch-clamp recording under the synaptic- and gap-junction-block drug cocktail (Fig. 2p and Extended Data Fig. 4k-n)^46^. We observed an increase in V_m_ and spike rate immediately after delivery of Na_2_SO_3_. In contrast, puffing either Na_2_SO_4_ (which is not an O_2_ scavenger) or extracellular solution (ES) caused no increase in V_m_ or spike rate (Fig. 2p,q and Extended Data Fig. 4m,n). To control for general effects of Na_2_SO_3_ on neurons that might not be restricted to those we found to be O_2_-sensitive, we tested the effect of puffing Na_2_SO_3_ onto the cerebellum while performing whole-cell recordings from these neurons (labeled as Cb in Fig. 2q,s). This produced no change in V_m_ or spike rate of cerebellar neurons (Fig. 2q and Extended Data Fig. 4m,n). Similar results were observed using calcium imaging (Fig. 2r,s). Together, these data show that, independently of peripheral sensors, NE-MO neurons can detect local changes in brain pO_2_.

### NE-MO neurons integrate brain pO_2_ and motor efference copies, consistent with predictive signaling for anticipated hypoxic stress

Hypoxia elevated the membrane excitability of NE-MO neurons and also increased fast calcium dynamics, which was coupled with swimming (Fig. 2c). Therefore, we hypothesized that brain pO_2_ would impact NE-MO spiking in response to motor signals. To test this, we performed simultaneous electrophysiological recordings of NE-MO and fictive swimming, and found a co-occurrence of excitatory postsynaptic potentials (EPSPs) in NE-MO and individual swim bouts (Fig. 3a), with swim bouts preceding the EPSPs by ~58 ms (Extended Data Fig. 5a), indicating that NE-MO cells receive excitatory motor efference copies. Consistent with increased fast calcium activity in hypoxia (Fig. 2c,e and Extended Data Fig. 5b-d), hypoxia led to an elevation of swim-coupled phasic spiking in NE-MO neurons (Fig. 3b–d). Therefore, the hypoxia-induced V_m_ rise appeared to make NE-MO cells more responsive to synaptic inputs, including motor efference copies (Fig. 3e and Extended Data Fig. 5h-k). We further tested this possibility by injecting current into NE-MO neurons to elevate their V_m_, which also led to an increase of swim-coupled spike rates (Extended Data Fig. 5e-g). Therefore, NE-MO can integrate current pO_2_ with motor efference copies, consistent with the predictive model combining both pO_2_ and swim history (Fig. 1j-l and Extended Data Fig. 1n,o). Through this approximately multiplicative interaction of hypoxia levels (~ 1/pO_2_) and motor signals via the membrane potential-to-spiking transformation (Fig. 3d,e), NE-MO spiking may represent predicted hypoxic stress, defined as the decrease in O_2_ levels relative to prior O_2_ availability or −ΔpO_2_/pO_2_, and thereby encode the urgency to preserve O_2_ under limited supply. Moreover, such a multiplicative interaction between hypoxia levels and swimming at NE-MO matched the prediction of the behavioral model (Fig. 1k). Remarkably, during normoxia, the slope of the approximately-linear relationship between swim vigor and NE-MO spiking was ~0.41 of that during hypoxia (Fig. 3d), closely matching the corresponding ratio of swim-history weights in the behavioral model (~0.44; Fig. 1k, inset). This concordance suggests that NE-MO activity implements the swim-history computation posited by the predictive model to transform current O_2_ availability and behavior into anticipated hypoxic stress.

**Figure 3.**
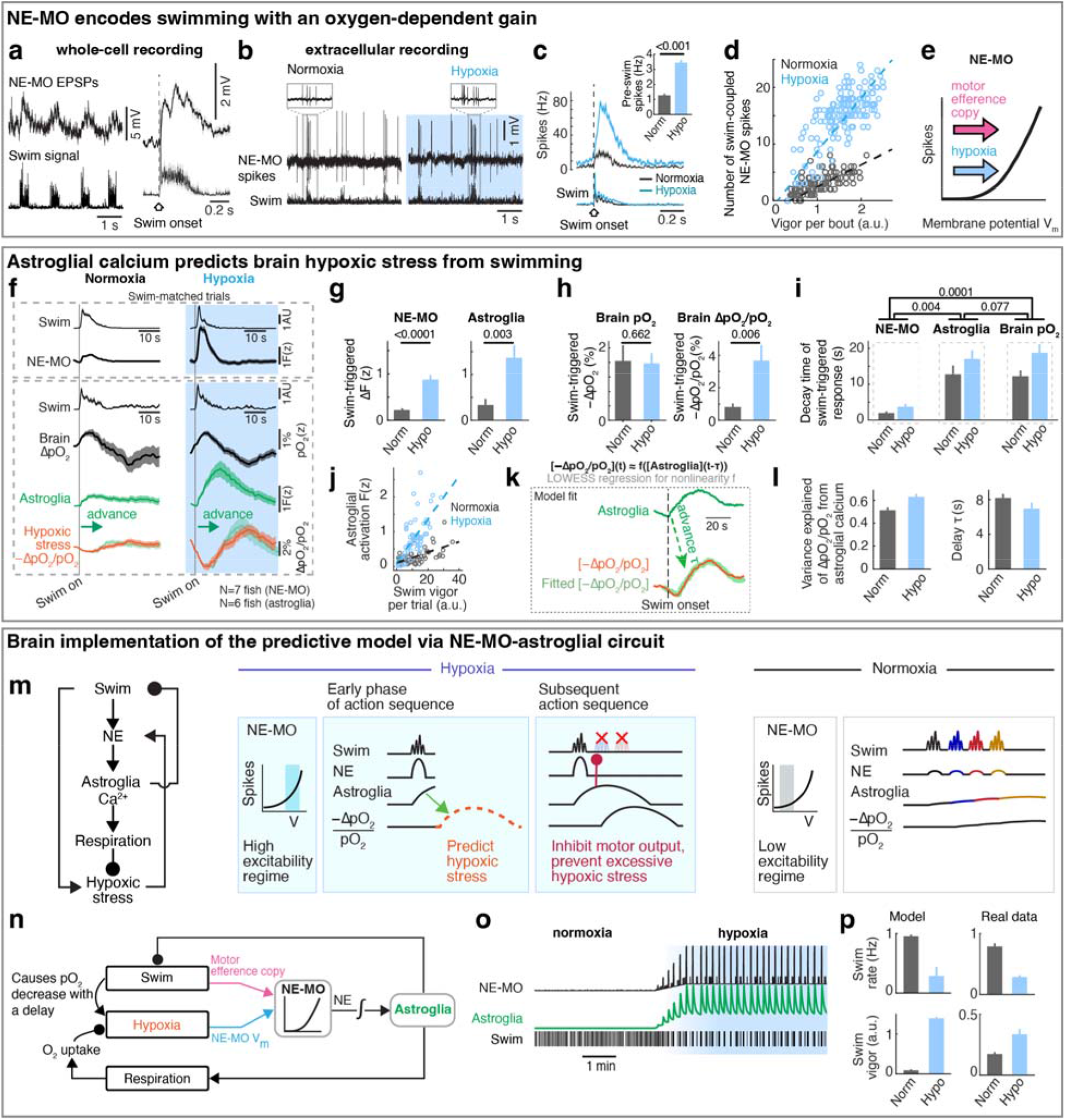
Predictive regulation of oxygen use by noradrenergic-astroglial signaling. **a**, Whole-cell recordings of NE-MO subthreshold voltage activity with simultaneous fictive swimming. Right, swim-triggered EPSPs for one cell across 60 swim bouts; mean ± s.e.m. is shown. **b**, Extracellular recording of a single NE-MO cell with simultaneous swim signals during normoxia (left) and hypoxia (right). Insets: zoom-in plots of electrophysiological recordings in the gray-boxed regions. **c**, Swim-triggered NE-MO spiking activity during normoxia and hypoxia. Bottom, averaged swim signal; inset, bar plots indicate mean ± s.e.m. of average pre-swim spikes; Wilcoxon rank-sum test. See also Extended Data Fig. 5c,d for comparison of swim-coupled NE-MO calcium activity in normoxia and hypoxia. **d**, Scatter plot of swim-coupled NE-MO spikes and swim vigor per bout during normoxia (black) and hypoxia (blue), implying an oxygen-dependent encoding of motor activity in NE-MO cells. Dashed lines indicate linear fits, reflecting a multiplicative operation: *[spike rate]=f(O*_*2*_*)×[swim vigor]*. **e**, Schematic of cellular model that explains the oxygen-dependent swim encoding in NE-MO cells. A NE-MO cell sums a hypoxia signal (which causes an increase in V_m_) and motor efference copies (via excitatory synaptic input) and passes these through an exponential nonlinearity to generate spikes. This model can account for *[spike rate]=f(O*_*2*_*)×[swim vigor]* in (d). See also the biophysiological implementation model in Extended Data Fig. 5h,i. **f**, Mean ± s.e.m. plots of swim-triggered dynamics of NE-MO (black), change in pO_2_ relative to baseline (black), astroglial calcium (green), and hypoxic-stress (pO_2_ decrease relative to pre-swim pO_2_; orange) during normoxia (left) and hypoxia (right), using swim-matched trials. Top box, simultaneously recorded swim and NE-MO dynamics; bottom box, simultaneously recorded swim, oxygen and astroglial dynamics. is shown. See also Video S1 for an animated description. **g**, Mean ± s.e.m. bar plots of swim-triggered calcium dynamics in NE-MO neurons (left) and astroglia (right) during normoxia and hypoxia. Calcium amplitudes were larger during hypoxia; Wilcoxon rank-sum test, across swim-matched trials in (f). **h**, Bar plots of swim-triggered changes in brain pO_2_ (left) and hypoxic stress (right) during normoxia and hypoxia. Swim-induced pO_2_ change was similar under hypoxia and normoxia, but hypoxic stress was higher during hypoxia due to normalization by pO_2_; Wilcoxon rank sum test. **i**, Mean ± s.e.m. of decay time constants of swim-triggered responses from NE-MO neurons (left), astroglia (middle), and brain pO_2_ (right). The decay time in both normoxia and hypoxia of NE-MO neurons is shorter than astroglia and pO_2_; that of astroglia is similar to pO_2_. Two-way ANOVA test based on the type of the dynamics and oxygen condition. **j**, Scatter plot of swim-triggered astroglial activation and swim vigor per trial (a swim series, lasting <40 seconds, flanked by pauses of >5 seconds) in (f) during normoxia (black) and hypoxia (blue); dashed lines indicate linear fits. **k**, Schematic of the model fit in which astroglial calcium at time (t − τ) predicts swim-induced hypoxic stress at time t, using LOWESS regression. The fit of astroglial calcium (green) and hypoxic-stress (orange) averaged for all trials for an example fish during normoxia. In this case, the delay τ is 4.3 seconds, variance explained is 0.98. **l**, Bar plots of the statistics of model fits (left) and astroglial-hypoxia delay (right) during normoxia (black) and hypoxia (blue). Mean ± s.e.m. across trials are shown. **m**, Cartoon illustrating predictive regulation of brain pO_2_ use via neuron-astroglial interactions. Left, model: swimming causes the increase of hypoxic stress; NE integrates hypoxia and swim actions; its output is then integrated in time by astroglial Ca^2+^; elevated astroglial Ca^2+^ inhibits swimming and promotes respiration, thereby alleviating hypoxic stress (see also Extended Data Figure 5m). Arrow, excitation; line ending in filled circle, inhibition. Middle, schematic showing time-series of changes during hypoxia. NE-MO encodes swim actions in a high excitability regime and thus predicts high hypoxic stress (green dashed line) via astroglia; the resulting large astroglial Ca^2+^ elevation then inhibits swim actions (red symbols) to prevent excessive hypoxic stress. Right, time-series of changes during normoxia. NE-MO encodes swim actions in a low excitability regime, wherein swims have only small effects on NE and astroglial Ca^2+^, resulting in only a small increase in hypoxic stress. As a result, subsequent swims are not suppressed. **n**, Schematic of the NE-MO–astroglial circuit implementing predictive regulation. Swimming induces delayed pO_2_ consumption. NE-MO neurons integrate motor efference copies with local brain pO_2_ signals through a multiplicative interaction (e), implementing predictions of hypoxic stress. Astroglia temporally integrate NE-MO spikes and suppress swimming while promoting oxygen uptake once predicted hypoxic stress exceeds a threshold. This circuit thereby preemptively pauses locomotion to prevent brain pO_2_ from falling below critical levels. Arrow, excitation; line ending in filled circle, inhibition. **o**, Computer simulation of the NE-MO–astroglial circuit (n) during transitions from normoxia to hypoxia. **p**, Comparison of hypoxia-induced changes in swim rate and swim vigor between experimental data (right; sampled from 29 fish) and simulation (left).

### Astroglial calcium is predictive of future hypoxic stress

The NE-MO representation of predicted hypoxic stress is transient (Fig. 3f, top), and thus insufficient to explain the prolonged integration of swim history in the predictive model (Fig. 1k). NE-MO spiking also does not, by itself, account for the intermittent, repeated ~18 second-long periods of locomotion suppression in hypoxia (Extended Data Fig. 1h), whose function may be to allow for sufficient time for pO_2_ recovery, which is also slow (Fig. 1h). We therefore tested whether astroglia, as a slow integrator of NE-MO activity^32^, could better predict the time course of future hypoxic stress from current swims. We examined swim-induced pO_2_ changes together with astroglial activity during sets of swims with similar vigor in normoxia and hypoxia (swim-matched trials) and found that astroglial calcium changes were similar in shape to the inverse of swim-induced O_2_ changes and, importantly, were shifted ~8 seconds earlier in time (Fig. 3f, bottom). Furthermore, swim events with similar vigor evoked larger astroglial calcium activity under hypoxia as compared to normoxia (Fig. 3g), consistent with the larger noradrenergic drive during hypoxia (Fig. 3c). Swim-induced pO_2_ decrease (−ΔpO_2_) was similar in the two conditions (Fig. 3h, left), but hypoxic stress (−ΔpO_2_/pO_2_) was larger during hypoxia due to the lower baseline level of pO_2_, mimicking astroglial scaling between the conditions (Fig. 3f and Fig. 3h, right). Moreover, hypoxic stress had a similar time course (Fig. 3i) and magnitude to astroglial dynamics (Fig. 3j). Similar to NE-MO (Fig. 3d), swim-triggered astroglial activation exhibited a hypoxia-dependent gain increase so that astroglial calcium signals encode a combination of swim vigor and depth of hypoxia, captured by the functional form F(z)~[1/pO_2_]×[integrated swim vigor] (Fig. 3j), suggesting both of them transform current O_2_ availability and behavior into anticipated hypoxic stress. Using LOWESS regression to compare astroglial and pO_2_ dynamics, we confirmed that, remarkably, astroglial activity accurately forecasted hypoxic stress for individual swims ~8 seconds in advance with high explained variances (0.51 in normoxia, 0.63 in hypoxia) (Fig. 3k,l). Moreover, optogenetic activation of astroglia in the hindbrain suppressed swimming and increased respiration (Extended Data Fig. 5l,m). These results show that astroglial calcium signaling is predictive of future hypoxic stress, i.e. predictive of pO_2_ decreases relative to pO_2_ availability. By encoding hypoxic stress several seconds in advance, and by its function of suppressing swim and promoting respiration, astroglial calcium signaling provides a time window to enable animals to preemptively adjust their behavior to minimize future internal hypoxia (Supplementary Video 1).

### Control theory accounts for neuron-glia dynamics

Based on NE-MO and astroglial dynamics predicting hypoxic stress (Fig. 3a-l), we extended the simple behavioral model (Fig. 1j) into a circuit implementation (Fig. 3m,n). This implementation was derived from a control model that optimized for a balance between brain pO_2_ and locomotion (Methods). This model recapitulated (1) the fish’s behavioral patterns and pO_2_ fluctuations during normoxia and hypoxia, (2) a multiplication of pO_2_ and swimming that resembles NE-MO activity, (3) an integration that resembles astroglial activity, and (4) occasional high vigor swims (reflected as an increase of swim bout duration both in simulation and real data) (Extended Data Fig. 1n,o). In simulation, this model recapitulated the change of swim patterns and noradrenergic-astroglial dynamics during normoxia and hypoxia (Fig. 3o,p). In summary, the optimal predictive model and its circuit implementation can account for the observed NE-MO-astroglial dynamics that underlies behavioral adaptation to hypoxia.

### A NE-MO-astroglial circuit mediates the hypoxia-induced brain and behavioral state

Since NE-MO shows the largest increases in both F_slow_ and ΔF/F in response to hypoxia, and can be directly activated by hypoxia, we asked whether the effects of hypoxia on brain state and behavior are dependent on NE-MO. We first ablated NE-MO using a two-photon laser in *Tg(th1:Gal4);Tg(UAS:NTR-mCherry)* fish, and found that the effects of hypoxia on neuronal and astroglial fast dynamics were almost completely abolished after NE-MO ablation (Fig. 4a-c and Extended Data Fig. 6a-g), while brain dynamics during normoxia were less affected. This result is consistent with a brain-wide modulatory role for NE. Specifically, hypoxia-induced functional relations among the neural clusters were suppressed by NE-MO ablation (Fig. 4d-f). For example, the normoxia and hypoxia ΔF/F manifolds of the midbrain-versus DRN-cluster and midbrain-versus forebrain-cluster were separated before NE-MO ablation, but became indistinguishable after ablation (Fig. 4e,f; see Methods). In summary, although neuronal slow dynamics retained some O_2_ dependence after the ablation (Extended Data Fig. 6b,f), potentially due to direct effects on cellular metabolism^39,51^, NE-MO mediated the hypoxia-induced modulations of brain-wide ΔF/F dynamics.

**Figure 4.**
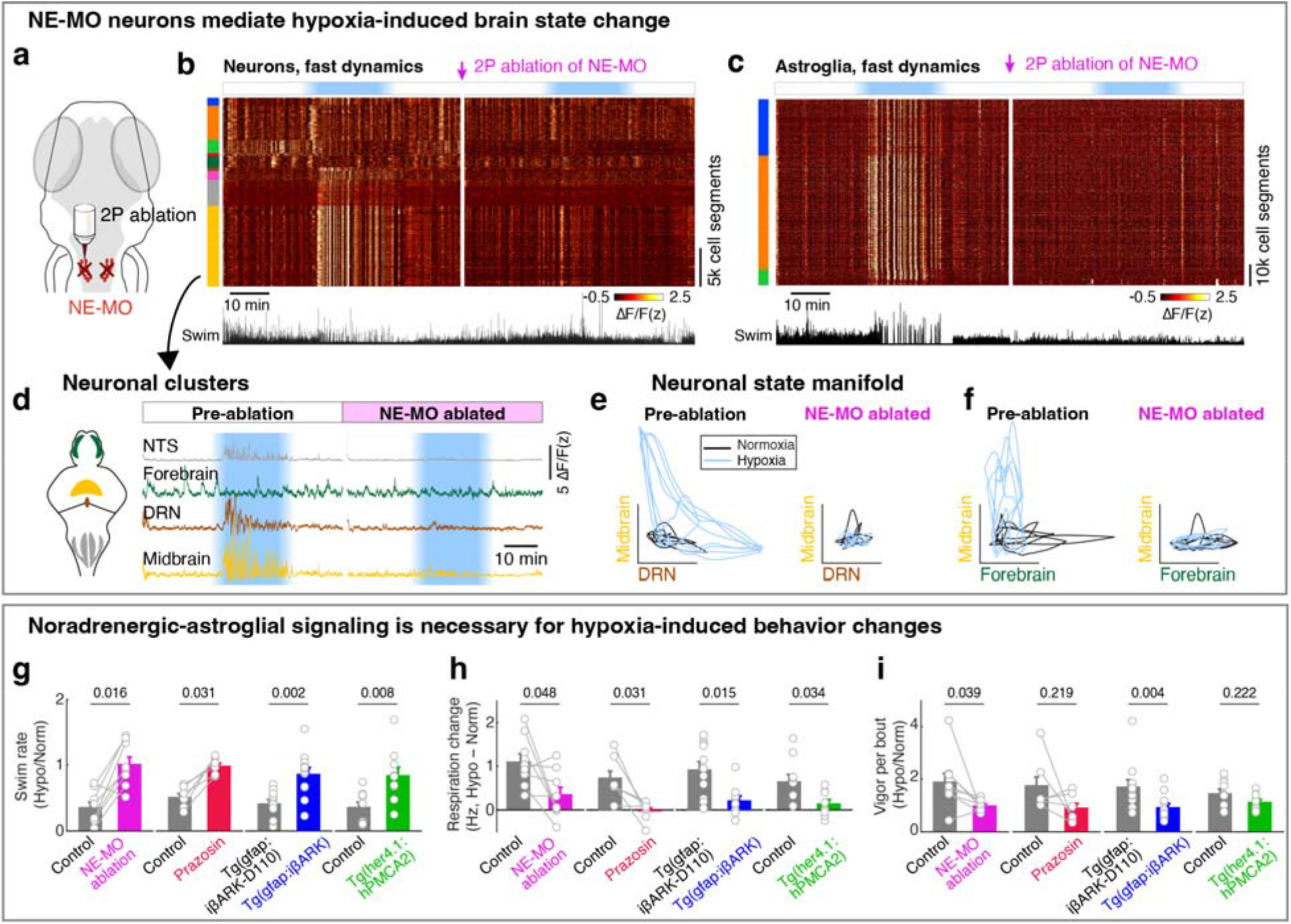
Noradrenergic-astroglial signaling is necessary for hypoxia-induced brain dynamics and coping behavior. **a**, Schematic showing two-photon laser ablation of NE-MO neurons. **b**,**c**, Example activity matrices showing the change of neuronal (b) and astroglial (c) cellular fast calcium (ΔF/F) responses to hypoxia before and after NE-MO ablation. Both matrices were clustered using factor analysis of the fast calcium component before NE-MO ablation. The spatial location and temporal responses for each cluster are shown in Extended Data Figure 6f,g. Bottom black traces show fictive swimming. **d**, Example traces showing calcium dynamics of 4 neuronal clusters before and after NE-MO ablation in the same fish. **e**,**f**, Overlaid embedding manifolds of neuronal cluster activity across 5 fish shows distinct dynamics under normoxia and hypoxia (left), which became indistinguishable after NE-MO ablation (right). Black and blue contours indicate the dynamics manifolds of the two clusters during normoxia and hypoxia, respectively. **i**, Effects of NE-MO ablation, prazosin treatment, or expression of iβARK or hPMCA2 in astroglia using *Tg(gfap:iβARK)* or *Tg(her4*.*1:hPMCA2)* fish on hypoxia-induced changes in swim rate (g), respiration rate (h) and swim vigor (i). *Tg(gfap:iβARK-D110)* fish express a mutant transgene that does not inhibit Gq GPCR signaling, and thus serves as a control. Dots indicate individual fish and bars indicate mean ± s.e.m. Wilcoxon signed-rank test for paired statistics in same groups with gray lines, Wilcoxon rank-sum test for unpaired statistics between groups. Due to low and variable respiration rate in these recorded fish during normoxia, we calculated the net change of respiration rate in (h). The effects of NE-MO or astroglial manipulation on behaviors during normoxia and hypoxia are also shown in Extended Data Figures 7-8.

In addition to suppressing the effect of hypoxia on brain state, ablation of NE-MO also substantially reduced hypoxia-induced changes of swim frequency, respiration rate, and swim vigor (Fig. 4g-i and Extended Data Fig. 7a). Conversely, mild and sustained optogenetic or chemogenetic NE-MO activation in normoxic conditions caused longer inter-swim-bout intervals (Extended Data Fig. 6h,i,l,m), as seen in hypoxia (Fig. 1e). Furthermore, optogenetically silencing NE-MO in hypoxic fish increased swimming and reduced respiration (Extended Data Fig. 6j,k). These NE-MO manipulations establish a causal link between NE-MO, brain-wide dynamics, and behavioral adaptations to hypoxia.

We next tested the role of noradrenergic signaling by blocking the NE-astroglia pathway with prazosin, a small molecule inhibitor of α1 adrenergic receptors (the predominant NE receptor on astroglia^52^). Treatment with prazosin almost completely abolished the effects of hypoxia on astroglial activity, swim frequency, respiration frequency, and swim vigor (Fig. 4g-i and Extended Data Fig. 7b,8a,8b). Notably, it also restored the optomotor response, which was otherwise inhibited during hypoxia (Extended Data Fig. 7e).

To explicitly test the role of astroglial activation in hypoxia, we suppressed the increase of astroglial Ca^2+^ that is normally triggered by NE using *Tg(gfap:iβARK)* or *Tg(her4*.*1:hPMCA2)* fish. In *Tg(gfap:iβARK)* fish, astroglia express an inhibitory peptide based on β-adrenergic receptor kinase 1 (iβARK)^53^ that disrupts intracellular signaling of Gq-coupled GPCRs, including α1 adrenergic receptors (Extended Data Fig. 8c). In *Tg(her4*.*1:hPMCA2)* fish, a calcium-transporting ATPase (hPMCA) is localized to the astroglial cell membrane, where it functions as a calcium extruder to reduce astroglial calcium levels (Extended Data Fig. 8i)^54,55^.

Consistent with the prazosin results, we found that hypoxia-induced changes in astroglial calcium responses, swim frequency, respiration rate, and swim vigor were almost completely abolished in *Tg(gfap:iβARK)* and *Tg(her4*.*1:hPMCA2)* fish (Fig. 4g-i and Extended Data Fig. 7c,d,8c-m). As a negative control, in *Tg(gfap:iβARK-D110)* fish, which express an inactive iβARK mutant protein that fails to disrupt Gq signaling, we observed normal astroglial calcium and behavioral responses to hypoxia (Fig. 4g-i and Extended Data Fig. 7c,8g,8h). Furthermore, whereas control fish almost stopped performing the optomotor response during hypoxia, this hypoxia-induced suppression of the optomotor response was largely prevented in *Tg(gfap:iβARK)* and *Tg(her4*.*1:hPMCA2)* fish: the astroglia-disabled fish behaved in hypoxia as they did in normoxia (Extended Data Fig. 7f,g). Therefore, noradrenergic and astroglial signaling are required for hypoxia-induced behaviors.

Taken together, these results reveal a neuromodulatory and astroglial predictive control system for adaptive behavior in the context of an essential but fluctuating metabolic resource.

## Discussion

To survive in variable environments, animals continuously regulate their physiology and behavior through reactive mechanisms (i.e. homeostasis) to correct for deviations in response to environmental or internal changes^7,56^. Examples of reactive regulation include sweating to maintain body temperature in hot weather, and drinking water after dehydration^57^. In addition to reactive homeostasis, animals also use anticipatory control (i.e. allostasis) to preempt physiological challenges^58^. For example, insulin is released upon tasting food, before nutrients enter the digestive system^59^, and ingestion of water causes rapid suppression of vasopressin release prior to the resulting change of blood osmolarity, thus preventing excessive renal water retention^60^. However, aside from a small number of well-studied cases, neural mechanisms for many forms of physiological regulation, especially predictive control, are poorly understood^3,58^. Here, we discovered that both behavior and brain dynamics of O_2_ regulation align with a predictive control strategy, through which future O_2_ consumption is estimated from swim history and used to proactively suppress swimming during resource scarcity. Our findings establish a mechanism of predictive O_2_ regulation, supported by convergent evidence across behavior, circuits, cells, and theory.

Although allostasis is often defined as predictive regulation^4,5^, it also has been framed as regulation via context-dependent set points^61^. Consistent with the predictive view, astroglial calcium is predictive of future hypoxia. Moreover, astroglial calcium integrates recent motor output scaled by recently experienced hypoxia to suppress future swimming, and this scaling is analogous to a context-dependent shift in set point for O_2_ preservation. Thus, our findings bridge these complementary views of allostatic control at the mechanistic level.

As animals encounter changing environmental conditions and physiological states, their behavior and physiology must adapt^9,62^. Despite a limited capacity for anaerobic metabolism, hypoxia is a threat to all cells of an animal and requires extensive behavioral adaptations and a global reconfiguration of brain dynamics. Neuromodulatory systems are well-suited to orchestrate such changes because they project across large swaths of the brain and can influence global brain dynamics^63,64^, sometimes through astrocytes^43,65–70^, which aligns with our discovery that a noradrenergic-astroglial system encodes and integrates hypoxic stress over ethologically-relevant long timescales to influence brain and behavior states. Moreover, this neuromodulatory-astroglial system can induce effects at different timescales, i.e. promoting vigorous swims at a short time scale via NE release^71^ and suppressing swimming via activation of astroglia at a longer time scale^32^, which agrees with the swim-pause cycles in hypoxia and biphasic swim kernel captured by the predictive model. Thus, this predictive biological system can provide proactive regulation that optimizes metabolic usage for behaviors during environmental and physiological challenges like hypoxia and futility^32^. Similar to hypoxic stress, peripheral sympathetic nerve terminals and central noradrenergic LC neurons can release norepinephrine in response to other stress stimuli, and orchestrate body-wide vigilance states and classic ‘fight-or-flight’ responses^72–74^. We found that noradrenergic neurons in the sympathetic ganglion and LC showed modest activation in hypoxia, suggesting that these noradrenergic populations may also contribute to hypoxia-induced physiological adaptations in synergy with noradrenergic neurons in NTS^75,76^. Considering their sensitivity to stimuli and widespread projections, noradrenergic neurons may act as allostatic mediators, capable of predicting impending challenges and orchestrating body-wide adjustments that optimize cardiovascular, metabolic, and behavioral responses to stressors^74^.

Relative to peripheral chemoreceptors, such as the mammalian carotid bodies, central neurons for O_2_ sensing and behavior execution are less well-studied^77,78^. Consistent with our results, catecholaminergic neurons and sympathetic nerves have been shown to respond to hypoxia and hypercapnia^79–81^. However, their molecular mechanisms for sensing O_2_, and the circuit connectivity underlying hypoxia-triggered behaviors, were largely unknown. The latter includes the neuron-glia coupling within breathing circuits that produce rhythmic activity^82,83^; these circuits have been mapped out at the regional scale in mammals and include the pre-Bötzinger complex^84,85^, but whether they directly translate to zebrafish is unknown. Ventilation in mammals increases immediately upon the onset of exercise—before any measurable changes in arterial O_2_, CO_2_, or pH occur—and carotid body chemoreceptors are reset at exercise initiation^86,87^, which exemplifies a predictive regulation of O_2_ demands similar to the strategy identified in our study. Additionally, land-roaming mammals exist in conditions that are less susceptible to large fluctuations in environmental O_2_ levels compared to fish, although burrowing mammals and those at high altitudes can encounter similar challenges^20,21,88^, choking and strangulation can cause hypoxia^22^, and diving or overexertion can also cause mammals to experience pO_2_ decreases resulting from their own body’s consumption, each of which requires behavioral adaptations^15,16,89,90^. Taking into account these commonalities and differences, we expect that core elements of allostasis are conserved across species; identifying specific similarities and differences will require additional studies. A second area of future investigation includes the molecular pathway underlying brain pO_2_ sensing, which is not fully understood even in peripheral chemoreceptors^48^. Possible mechanisms include local neuron-glial interactions as seen in the retrotrapezoid nucleus area of mammals^82,83,50,91^, reactive oxygen species (ROS) production in cells, and ion channel modulation by metabolic products such as ROS, ATP, or lactate^48^, but the full space of possibilities requires exploration.

## Methods

### Zebrafish

Larval zebrafish were reared in system water at 28.5°C with 14:10 hour light:dark cycles. Zebrafish from 6 to 9 dpf were fed with rotifers and used for experiments. At this stage of development, sex is not defined. Transgenic zebrafish larvae were in a *casper* or *nacre* mutant background, and were generated using the Tol2 system^92^, except that *Tg(th1:gal4)* was generated by CRISPR/Cas9-based knock-in^93^. All experiments were conducted according to the animal research guidelines from NIH and approved by the Institutional Care and Use Committee (IACUC) at Janelia Research Campus (animal protocol 22-0216) and IACUC at California Institute of Technology (animal protocol 1836).

### Transgenesis

For imaging neuronal or astroglial activity we used transgenic zebrafish lines expressing:

- Nuclear-localized jGCaMP7f in all neurons, *Tg(elavl3:H2B-jGCaMP7f)*^94^
- Cytosolic jRGECO1b in all neurons, *Tg(elavl3:jRGECO1b)*^95^
- Cytosolic GCaMP6f in all astroglia, *Tg(gfap:GCaMP6f)*^32^
- Cytosolic jRGECO1b in all astroglia, *Tg(gfap:jRGECO1b)*^32^
- Cytosolic GCaMP6f in all noradrenergic neurons, *Tg(th1:gal4); Tg(UAS:GCaMP6f)*^93^
- Cytosolic GCaMP8f in peripheral ganglia, *Tg(17768:gal4);Tg(UAS:jGCaMP8f)* (this paper)

For norepinephrine release imaging, we used:

- *Tg(elavl3:GRAB-NE)* (this paper) For silencing astroglial activity we used:
- *Tg(gfap:iβARK)* (this paper)
- *Tg(gfap:iβARK-D110)* (this paper)
- *Tg(her4*.*1:PMCA2)* (this paper)

For optogenetic stimulation, optogenetic inhibition, and chemogenetic stimulation we used:

- *Tg(gfap:CoChR-eGFP)*^32^
- *Tg(th1:gal4); Tg(UAS:GtACR2-eYFP)* (this paper)
- *Tg(dbh:KalTA4); Tg(UAS:CoChR-eGFP)*^32^
- *Tg(dbh:KalTA4); Tg(UAS:TRPV1-mCherry)* (this paper)

For 2-photon ablation and whole-cell patch-clamp recording we used:

- *Tg(th1:gal4); Tg(UAS:NTR-mCherry)* (this paper)

### Preparation of oxygenated solutions

Zebrafish larvae were maintained in zebrafish facility system water, whose dissolved O_2_ concentration was ~8.5 mg/L (i.e. 150 mmHg pO_2_), which was measured using a portable O_2_ meter (MW600, Milwaukee). To generate water with low or high dissolved O_2_ levels, system water was bubbled with nitrogen or oxygen gas for 30 minutes prior to each experiment. The resulting nitrogen- or oxygen-bubbled water was then mixed with untreated system water to obtain a series of intermediate O_2_ concentrations ranging between 1 and 20 mg/L (i.e. between 20 and 350 mmHg). The final dissolved O_2_ concentration of each mixture was measured using an O_2_ meter. In most experiments, system water with 4 mg/L or 16 mg/L was used for hypoxic or hyperoxic treatment, respectively. pH, measured using a pH meter (SD20, Mettler Toledo), was the same regardless of O_2_ content.

### Fictive swimming and fictive respiration

Zebrafish larvae were paralyzed with alpha-bungarotoxin (1 mg/ml, Invitrogen), then embedded in 1.5% low melting-point agarose (Sigma) with dorsal side up in a custom-made chamber, as previously described^96^. Agarose was removed above the head and around the tail to expose those regions for electrophysiological recordings. Microelectrodes for recording fictive swimming and fictive respiration were of the same size, made with borosilicate capillaries (TW150-3, World Precision Instruments), pulled by a vertical puller (PC-10, Narishige, Japan) and polished by a microforge (MF-900, Narishige, Japan) to make pipettes with a tip diameter of ~50 μm. Extracellular recordings of fictive swims were made by attaching a pipette to the tail with slight suction, as previously described^94^. Electrophysiological signals were captured using an Axon Multiclamp 700B amplifier in current clamp and acquired using National Instruments DAQ boards (PCIe-6321, BNC-2110), sampled at 6 kHz, band-pass filtered with a 300 Hz/1000 Hz high-pass/low-pass cutoff for swim signals, and recorded by custom software written in C# (Microsoft) which is available upon request. Individual swim bouts and swim vigor were detected offline using a custom Matlab script as previously described^96^. In brief, raw electrical signals representing fictive swimming were processed using a windowed (10 ms) standard deviation to generate a smoothed power of swim signal. A histogram distribution of power was used to determine a threshold, which was then used for detecting the start and end of each swim bout when the amplitude of the swim signal was above the threshold. The integral from the start to the end of one swim bout was computed as the swim vigor of that bout. Only active fish with swim frequency larger than 0.4 Hz under normoxia were included in the analysis.

Extracellular recordings of fictive respiration were made by placing a pipette onto the skin above the optic tectum or the tail, with slight suction to obtain a good seal. In the preparation, respiration-related electrical activity could be detected by the large tip-opening pipettes. The recording configuration was the same as that for fictive swims, except that it was filtered with a 0.1-100 Hz band-pass filter. After the experiments, the fictive respiration signal was further processed with a 10 Hz low-pass filter to remove high-frequency noise. The fictive respiration events were detected by tracking the peak amplitude of electrical respiratory activities during the periods between swim bouts using the “findpeak” function in MATLAB. Extended Data Figure 1a-c validates this approach for recording fictive respiration by simultaneous detection of respiratory behaviors.

### Light-sheet imaging

All whole-brain imaging experiments were performed with a custom light-sheet microscope as previously described^97^. Briefly, two low-magnification objectives (4x/0.28 NA, Olympus) were used to generate excitation laser beams, which were scanned in the horizontal and vertical planes and controlled by galvanometer scanners (Cambridge Technology). Larval zebrafish were placed in a custom-made chamber with glass windows, so that the laser beams were focused through the glass windows onto the fish brain. The side laser was turned off when it was pointed at the eye to avoid direct stimulation of the retina. The detection arm consists of a water-dipping objective (16x/0.8 NA, Nikon), a tube lens, and an sCMOS camera (ORCA Flash 4.0, Hamamatsu). In general, whole-brain neuronal and astroglial volumetric images were acquired at ~2-3 volumes/second, with 5 or 10 μm between imaged planes. Blue (488 nm) and green (561 nm) lasers (SOLE-3, Omicron) were used to excite GCaMP and jRGECO1b, respectively, and green 525/50 nm or red 630/92 nm filters (Semrock) were used for detection. For two-color imaging, the 488 nm and 561 nm lasers were used simultaneously, and an image splitter (W-View Gemini, Hamamatsu) was used to separate the green and red images. The data acquisition was based on custom software written in LabVIEW, as described previously^94,97^.

For visual stimulation, a stripe pattern (2 mm stripe width) was projected underneath the fish chamber by a video projector (WT50, Akaso). A red Kodak filter (Wratten #25) was used for visual stimulation when imaging jGCaMP7f, whereas a blue Kodak filter (Wratten #47) was used for visual stimulation when imaging jRGECO1b or when imaging both jGCaMP7f and jRGECO1b.

Fish were placed in a virtual-reality (VR) system. In open-loop VR mode, the red/black visual stripes moved forward at a fixed speed of ~2.5 mm/s. In closed-loop VR mode, the visual stripes moved at 2.5 mm/s when the fish was not swimming, and additionally moved backward at a rate proportional to swim vigor when the fish was swimming. This mimics the visual feedback the animal would receive in a freely-swimming environment. In spontaneous VR mode, the fish stayed in a light background with red and black stripes. In each of these environments, relative to normoxia, hypoxia caused decreases in swim frequency, increases in respiration frequency, and increases in NE-MO and astroglial calcium activities.

### Whole-cell patch-clamp recordings

Fish were paralyzed as described for whole-brain imaging and extracellular electrophysiological recordings. After paralysis, fish were embedded in a custom-made glass-bottom Petri dish, and bathed with extracellular solution, which consists of (in mM) 134 NaCl, 2.9 KCl, 2.1 CaCl_2_, 1.2 MgCl_2_, 10 HEPES and 10 glucose (290 mOsmol L^−1^, pH 7.8). A small incision was made in the skin above the ventricle between cerebellum and hindbrain using a glass micropipette with a tip opening of 1 μm as previously described^98^. Micropipettes were made from borosilicate glass (BF100-58-10, Sutter Instrument) by a Flaming/Brown micropipette puller P-97 (Sutter Instrument). The prepared fish was then transferred to an electrophysiological recording set-up, and perfused with extracellular solution at ~4 ml/min using a peristaltic pump (Longer Pump). *In vivo* whole-cell patch-clamp recordings were performed using micropipettes with resistance of 20-30 MΩ filled with internal solution containing (in mM) 100 K-gluconate, 10 KCl, 2 CaCl_2_, 2 Mg-ATP, 2 Na_2_-GFP, 10 HEPES, and 10 EGTA (280 mOsmol L^−1^, pH 7.4). Neurons in the LC, MO, AP and cerebellum were visualized using *Tg(th1:Gal4); Tg(UAS:NTR-mCherry)* fish under fluorescent signal guidance using an upright microscope (BX51WI, Olympus). Contact of the micropipette onto neurons was confirmed by visualizing mCherry-positive membranes within the tip of the recording micropipette. An EPC-10 amplifier and patchmaster software (Heka, Germany) were used for electrophysiology and signals were filtered at 2.9 kHz and sampled at 10 kHz.

Spontaneous neuronal activity of most cells was recorded under current clamp (I=0 pA) without stimulus, while some data were captured via extracellular unit recordings. The data were accepted if the series resistance was below 100 MΩ, and the baseline of resting membrane potential or spike firing frequency was stable for more than 10 minutes before perfusion of hypoxic water.

The analysis of electrophysiological data and spike detection were processed using Clampfit (Axon instrument) and Matlab. We set a maximal inter-spike interval of 70 ms for phasic spike firing of noradrenergic neurons. The minimum number of spikes in a burst was set at 3. We analyzed total spike rate, tonic spike rate and phasic burst events per minute for Fig. 2 and Extended Data Fig. 4.

### Measurement of brain and bath O_2_ levels

During imaging and electrophysiological assays, the recording chamber was continuously perfused with system water at a speed of ~4-5 ml per minute. The water inflow was controlled by gravity and valves, and the outflow was controlled by vacuum suction. O_2_ levels in the brain and in the bath solution were recorded with calibrated Clark-type polarographic O_2_ microelectrodes (OX-10, Unisense A/S)^27,99^ with a customized tip diameter of 3-5 μm, whose 90% response time is ~0.3 s (T_90_, the time for 90% of final measured signal is reached). The O_2_ probe for brain pO_2_ was inserted into the anterior hindbrain at a depth of ~50-100 μm using a manual manipulator (MX130, Sisikiyou), and the O_2_ probe for bath O_2_ levels was placed in the bath solution. These O_2_ signals were amplified by a high-impedance amperometric device (fx-6 UniAmp, Unisense A/S) and converted by an analog-to-digital device (BNC-2110 and PCIe-6321 board, National Instruments) with the same configuration as for electrophysiological recordings. The data were denoised using a 10 second window using the ‘smooth’ function in MATLAB and analyzed offline.

### Puffing of O_2_ scavenger

In O_2_ scavenger experiments, sodium sulfite (Na_2_SO_3_, O_2_ scavenger) or sodium sulfate (Na_2_SO_4_, control) was dissolved in extracellular solution, loaded into glass micropipettes, and ejected using a PicoSpritzer III (Parker Instruments). For electrophysiological experiments, we used ~2-3 μm tip diameter pipettes and ~5 psi air pressure for 0.5 seconds at 30-second intervals. For imaging experiments, we used ~5 μm tip diameter pipettes and ~10 psi air pressure for 5 seconds at 120-second intervals in order to eject more solution around the tissue. In Fig. 2p,q we calculated the change of spike firing by dividing the values 2 seconds after the start of puffing by the baseline before puffing. In Extended Data Fig. 4m,n we calculated the change of V_m_ by subtracting the values 2 seconds after the start of puffing from the baseline before puffing.

To examine the effect of puffing O_2_ scavenger on pO_2_ change in Extended Data Fig. 4k,l, a ~5 μm tip diameter pipette was placed ~20 μm away from the tip of the O_2_ probe. Different concentrations (0.01-1.0 M) of Na_2_SO_3_, Na_2_SO_4_ or external solution were used for the calibration. The Unisense O_2_ probe OX-10 is sensitive to shear force. Therefore, puffing the control solution (external solution or 20 mM Na_2_SO_4_) caused a positive deflection of pO_2_ value. In comparison, puffing 20 mM Na_2_SO_3_ caused a smaller positive response. Therefore we set the control puffing solution value (~ +0.67 mg/L) as the baseline, and the net pO_2_ change by puffing 20 mM Na_2_SO_3_ was about −0.4 mg/L. Notably, puffing 100 mM or 1 M Na_2_SO_3_ caused negative responses, similar to published results in mouse somatosensory cortex^28^. We found that puffing high concentrations (100 mM or 1 M) of Na_2_SO_3_ resulted in a large calcium response, followed by a lack of responses to additional puffs. This was not an issue with puffs of 20 mM Na_2_SO_3_ or Na_2_SO_4_, so we used this concentration for the electrophysiological and imaging experiments in Fig. 2p-s.

### Pharmacology

A drug cocktail including D(−)-2-Amino-5-phosphonopentanoic acid (APV, 50 μM), 6-cyano-7-nitroquinoxaline-2,3-dione (CNQX, 50 μM), hexamethonium (HEX, 100 mM), atropine (2 μM), carbenoxolone (CBX, 100 μM) was used to block conventional NMDA and AMPA receptors, nicotinic and muscarinic acetylcholine receptors, and gap junctions in synaptic-block and O_2_ scavenger experiments (Fig. 2k-s and Extended Data Fig. 4g-n). Tricaine (0.02%) was used to block sodium channels (Extended Data Fig. 2g,h).

### 2-photon neuronal ablation

To ablate NE-MO cells (Fig. 4 and Extended Data Fig. 6a-g), we used a two-photon titanium:sapphire laser (Chameleon Ultra II, Coherent) as previously described^94^. High laser power (~2500 mW at the laser at 90% maximum) at 850 nm with a short exposure time (5 ms) was applied onto cell bodies of fluorescent NE-MO cells using *Tg(th1:gal4); Tg(UAS:NTR-mCherry)* fish at a 500 ms interval. Ablations were confirmed one hour later. Fish swimming had usually recovered by that time, and a second hypoxia treatment was applied to determine the effect of NEMO ablation on brain activity and behavior.

Nodose ganglion and olfactory epithelial cells were ablated (Extended Data Fig. 3) using *Tg(elavl3:H2B-jGCaMP7f)* or *Tg(elavl3:jRGECO1b); Tg(gfap:GCaMP6f)* fish the night before an experiment using a Zeiss 880 microscope equipped with a two-photon laser (Chameleon Ultra II, Coherent) and 20x water immersion objective (1.0 NA). To ablate the nodose ganglion, fish were embedded in agarose on their side to ablate one side. The fish was then released and embedded again to ablate the nodose ganglion on the other side. The ablation was considered successful when no GFP was observable and a small cavitation was briefly visible after laser treatment. Only healthy fish with normal swimming patterns were used for experiments on the next day.

### DMD-based optogenetics

To specifically activate NE-MO or astroglial cells in the hindbrain, we applied optogenetic stimulation using a digital micromirror device (DMD) as previously described^32^. The DMD (LightCrafter DLP4500, Texas Instruments) was illuminated by a blue laser (488 nm, SOLE-3, Omicron). The patterns for target stimulation areas were generated using a custom Python script, and applied to the DMD using custom software written in C# (Microsoft). To mimic fast calcium increases of astroglial cells under hypoxia, we applied a series of 10-ms pulse illuminations at 10-Hz for 5 seconds followed by 25-s light-off periods. The laser intensity measured under the objective was ~0.15 mW/mm^2^ for experiments using *Tg(gfap:CoChR-eGFP)* fish in Extended Data Fig. 5l,m. To mimic slow activity changes of NE-MO under hypoxia or hyperoxia, we applied a continuous (60 seconds), but low-intensity light onto the target cells followed by a 60-s light-off period. The laser intensity measured under the objective was ~0.06 mW/mm^2^ for experiments using *Tg(dbh:KalTA4); Tg(UAS:CoChR-eGFP)* fish in Extended Data Fig. 6h,i; and ~0.03 mW/mm^2^ using *Tg(th1:gal4); Tg(UAS:GtACR2-eYFP)* fish in Extended Data Fig. 6j,k.

### Cell segmentation

We extracted populations of cell bodies and neuropil from raw fluorescence data using a previously described volumetric segmentation pipeline^94^ that is available at https://github.com/mikarubi/voluseg.

### Anatomical brain atlas preparation

We constructed a neuronal brain atlas by imaging *Tg(elavl3:H2B-jGCaMP7f)* fish at 0.406 μm x 0.406 μm x 2 μm (x, y, z; 2509 × 1325 × 183 voxels) resolution, and the astroglial brain atlas by imaging *Tg(gfap:GCaMP6f)* fish at 0.296 μm x 0.296 μm x 1 μm (x, y, z; 3200 × 1700 × 281 voxels) resolution using a one-photon confocal microscope (LSM 980, Zeiss).

### Brain registration

We registered the brain images of *Tg(elavl3:H2B-jGCaMP7f)* and *Tg(elavl3:GRAB-NE)* to the neural brain atlas and *Tg(gfap:GCaMP6f)* fish to the astroglial brain atlas using the “greedy” algorithm^100^. The details of the algorithm can be found in the github repo at https://github.com/zqwei/Hypoxia_dynamics.

### Fast-slow dynamics decomposition

We found that slow dynamics on timescales of ~5 minutes represents the trend of the internal brain oxygen (pO_2_), while fast dynamics represents fish behaviors like swim vigor. We thus decomposed the fluorescence dynamics F (raw fluorescence) of cell segments as

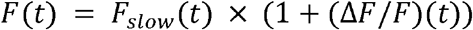

where *F*_*slow*_ is the slow dynamics computed as the 20^th^ percentile in a 5-minute window centered at each time point, and Δ*F*/*F* is the fast dynamics.

### Factor-based whole brain clusters analysis

We ran a sparse version of factor analysis^101^ onto whole brain data. In brief, we reconstructed the neural data matrix

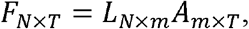

where *L*_*N* × *m*_ is the weight matrix, which reconstructs the neural dynamics from factors *A*_*m* × *T*_. *N* is the number of the cells, *T* is the number of time points, and *m* is the number of the factors. To increase the interpretability of the factors, we forced the sparsity of the weight matrix using a Beta distribution as a Bayesian prior distribution for the weights before sampling, *Beta*(0.5,0.5). We further separated the positive and negative weights in each factor into two factors.

### Factor combined across fish

We thresholded and binarized the weight matrices at 0.3 and combined the similar spatial factors across fish (Figs. 1m-q, and 4a-f, and Extended Data Figs. 2b-h, 3f-l, 6a-g, 8b-m). We counted the overlaps of the weight matrices across fish. The number of the fish with positive weight per voxel on the neuronal or astroglial brain atlas were computed within a diameter of 10 μm x 10 μm x 20 μm (x, y, z) per voxel.

### Slow and fast dynamics map

We constructed the slow dynamics map by calculating the correlation coefficient between slow response *F*_*slow*_ and the averaged brain pO_2_ dynamics (which were recorded independently using the O_2_ sensor in the same experimental paradigm) using rank correlation. We combined the positive and negative indices separately across fish (Fig. 1m-q and Extended Data Fig. 2b), which was averaged in a spatially moving spherical window of diameter 10 μm x 10 μm x 20 μm (x, y, z). Similarly, we constructed the normalized *F*_*slow*_ map by dividing the average of *F*_*slow*_ during hypoxia by that during normoxia (Fig. 2b and Extended Data Figs. 2h, 8b,f,h,k,m). We constructed the fast dynamics map by calculating the index d-prime, which is based on the comparison of the fast response Δ*F*/*F* in the last 5 minutes during the hypoxia epoch (from 35 to 40 minutes in our experiment paradigm) versus that in the last 5 minutes during the normoxia epoch (from 15 to 20 minutes in our experiment paradigm). We combined positive and negative pO_2_ modulation separately across fish (Fig. 1m-q and Extended Data Fig. 2c-f), which was averaged in a spatially moving spherical window of diameter 10 μm x 10 μm x 20 μm (x, y, z).

### Swim-matched event analysis

To examine our cellular model, we subsampled swim events (trials) with similar vigor. In these trials, we required that (1) the total vigor per swim is similar across O_2_ conditions; (2) the swim length is similar; (3) and the inter-swim-interval from the previous swim is longer than 1 second (so that the change of fluorescence is little affected by the previous swim). Trials were sampled in the late phase of normoxia (10 to 20 minutes) and hypoxia (30 to 40 minutes) conditions with a minimum of 10 swim events in each condition. The significance of the O_2_ condition was determined using the rank sum test on the average increase of fluorescence from the onset of swims in a 1 second time window.

### Circuit model

We formulated a circuit model (Fig. 3n-p) to mimic a variety of behavioral observations of swim patterns and fluorescence dynamics, Δ*F*/*F*(*t*), for NE-MO and astroglia. We simulated the ‘natural’ swim bouts from discrete ‘swim events’, where a swim bout may consist of one or multiple swim events. Each swim event is generated by a central pattern generator (CPG), whose output can be modulated by astrocytes or other neuromodulatory systems. This modulation adjusts the probability of swim-event generation, *P*_*swim*_, which can be down- or up-regulated. (In the below, *x* ′ (*t*) = *dx*(*t*)/*dt*.)

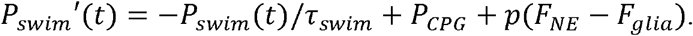

The underlying swim generator *P*_*CPG*_ - 1 every 1s (which generates swim bouts at 1 □ Hz in the absence of modulation); swim probability *P*_*swim*_ ∈ [0, 1] was clipped to this range; the swim modulator *p*(*x*) = *x*/3 is clipped to [0, 1] and depends on the level of NE (*F*_*NE*_) and glia (*F*_*plio*_). A swim event was generated when *P*_*swim*_(*t*) > *P*(*t*), where *P*(*t*) is a random number sampled uniformly from (0,1). *τ* _*swim*_ = 50 ms (such that a swim event lasts for ~100 ms) approximates the half-decay time of a swim bout containing a single swim event in real data.

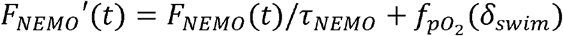

where *τ*_*NEMO*_ = 100 ms, 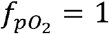 for normoxia and 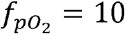 for hypoxia.

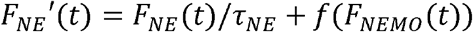

where *τ*_*NE*_ = 100 ms, *f*(*x*) = *RELU*(*x* — *x*_0_), *x*_0_ - 0.3.

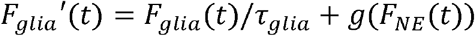

where *τ*_*plio*_ = 3 s, *g*(*x*) = *gx, g*= 3. When the CPG was sufficiently excited to produce multiple swim events with inter-event intervals <50 □ ms, these events were grouped into a single swim bout; otherwise, isolated swim events were treated as individual bouts. To enable comparison with experimental data (Fig. 3p), swim vigor was quantified as the number of swim events within each swim bout.

### A delayed control system – a normative model framework for oxygen allostasis

We assumed that the brain oxygen dynamics, *x*(*t*), was updated by oxygen uptake from environment, *x*^*e*^, and oxygen consumption from swimming, *u*(*t*). Since the oxygen consumption went possibly through a cascade process, from spinal cord motor neural activity to muscle contraction to tissue and blood oxygen depletion^38^, we simplified this process as a system delay, *τ*. Therefore, the normative model for this delayed system would be:

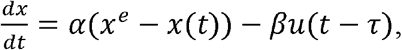

where *α* is the oxygen uptake rate; *β* is the oxygen consumption rate.

Here the system minimizes the infinite horizon cost functional, J(x,u),

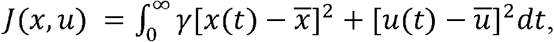

where 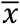 is the desired oxygen level of the system; ū is an averaged swim rate generated by swim CPG; γ is the weight between the two components of the objective function. The second term in optimization is to maintain the overall swim rates – fish tend to not swim at excessively high rates, nor should they stop swimming altogether since that would presumably be fatal, whether in normoxia or hypoxia.

We derived the linear optimal delayed feedback controller^102,103^ as

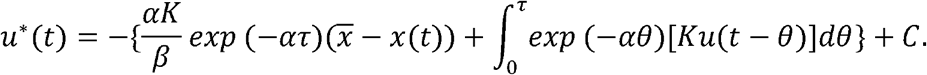

Here the gain 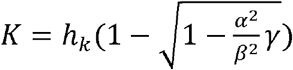 and 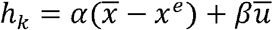, which predicts: (1) during normoxia, this K is small 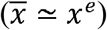 and the integral term is less effective in controller; (2) during hypoxia, K produces a notable multiplicative effect of hypoxic level and swimming signal as 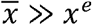. The constant 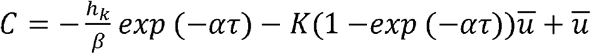 represents a ‘natural’ rate of swimming, which decreases as *x*^*e*^ decreases.

The model can be mapped into two processes: the first term, 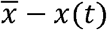, is represented as a reactive control from the slow dynamics of the measured oxygen. The second term reflects predictive control that integrates swims to predict future oxygen consumption. The term *Ku*(*t* – *θ*) reflects oxygen-dependent swim vigor (Fig. 3d). The overall modulation, i.e. slow dynamics in the first term plus fast dynamics in the second term, would inhibit future swimming (represented as the minus sign).

### Model fits

We fitted a generalized linear model of behavioral data (Fig. 1i-l), binned by 1 second. This model predicts the probability of a swim action (1, swim; 0, no swim) at time *t* based on a swim history from *t-9* to *t-1* and oxygen history from *t-8* to *t* as follow:

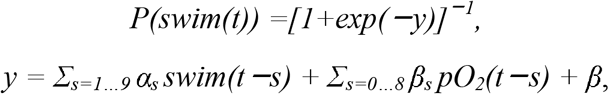

where *swim(t)* is the swim action, *pO*_*2*_*(t)* is the brain pO_2_ at time *t*; *α*_*s*_, *β*_*s*_ are the swim and oxygen kernels, respectively; *β* is the offset term. We fitted this model of the behavioral data under normoxia and hypoxia separately, where the offset *β* could represent the external oxygen condition. Goodness of fit was measured using pseudo variance explained^104^, *r*^*2*^ *= 1* − *ln(L*_*M*_*)/ln(L*_*0*_), where *L*_*M*_ is the likelihood of the fitted model and *L*_*0*_ is the likelihood of the null model (model fitted only by the offset *β*). We performed the model fits with both swim and oxygen history to quantify the variance explained by the predictive model (Fig. 1j) and performed the model fits with only oxygen history to quantify the variance explained by the reactive model (Fig. 1i).

We applied the LOWESS regression^105^, a locally linear regression, to predict hypoxic stress (Fig. 3k,l), −ΔpO_2_(t)/pO_2_(t), at time t from the astroglial dynamics, astroglia(t−τ), at time t−τ. Here we estimated hypoxic stress as

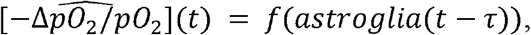

where the nonlinearity f is determined by the LOWESS regression and the optimal τ is determined by comparing the performance of the fits from a grid search of τ. Goodness of fits were measured using variance explained,

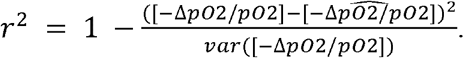

For all model fits, we used default setups and hyperparameters from the Python package Statsmodels^106^.

### Statistical analysis

Graphs show mean ± s.e.m. unless stated otherwise. Statistical significance in most cases was determined using nonparametric statistical tests: the Wilcoxon signed-rank test for paired data, and the Wilcoxon rank-sum test for independent samples. Student’s t-test was used if the data was normally distributed, which was determined using the Shaprio-Wilk test. Rank correlation was used for correlation analysis. No statistical methods were used to predetermine sample size.

## Supporting information

Video S1: Animated description of Figure 3f

## Acknowledgements

We thank Emmanuel Marquez Legorreta for helping to prepare and image the confocal astroglial reference brain. We thank Ann Hermundstad, Yin Liu, Grigorios Oikonomou, Dhruv Zocchi, and Anoj Ilanges for comments on the manuscript. We thank Janelia Experimental Technology for creating the behavioral arena, and the Vivarium staff for fish care. We thank members of the Du, Ahrens, and Prober labs for assisting with experiments and idea formation. We thank Joshua Vogelstein for intellectual input. We thank the Janelia Visiting Scientist Program for support. This study was supported by grants from the National Institutes of Health (R35 NS122172 D.A.P.) and the Howard Hughes Medical Institute (M.B.A.).

## Author contributions

RZ, ZW, DAP, MBA conceived the project, analyzed the data and wrote the manuscript. RZ performed imaging, behavior, and electrophysiology experiments, and data analysis for behavior, neural dynamics, astroglial dynamics and noradrenergic dynamics. ZW developed and performed population analyses for neural dynamics, developed models for optimal control, network dynamics, and biophysics, and developed data processing methodology. JJH developed the behavioral analysis pipeline. MN derived the brain-body control system model. SN and J-XL developed transgenic fish. AK assisted with behavioral analysis. WC helped with data preprocessing and managed fish lines. VMSR and BDM contributed intellectual input. AR and BB provided support for biophysical model conceptualization. MH and FE provided *Tg(17768:gal4)* fish. MCF provided support in data conceptualization and interpretation. JD provided support for electrophysiological experiments.

## Additional Information

Supplementary Information is available for this paper.

Correspondence and requests for materials should be addressed to.

## Extended Data Figures

**Extended Data Figure 1.**
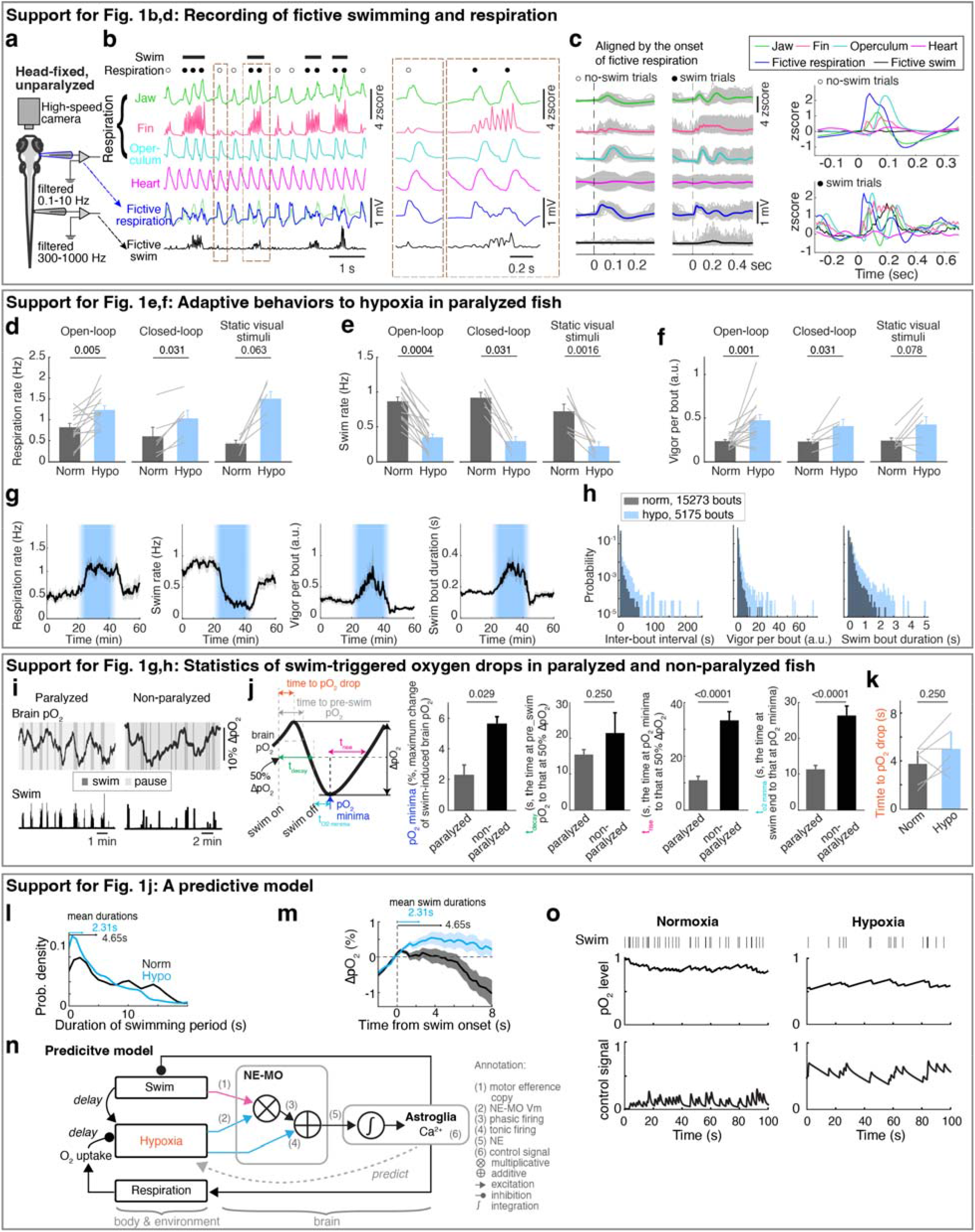
Oxygen-dependent behavioral adaptations and the predictive model. **a**, Schematic of simultaneous bright-field imaging of craniofacial behavior and electrophysiological recordings of fictive respiration and swimming of an embedded, but non-paralyzed larval zebrafish. A non-invasive electrophysiological assay is used to record fictive respiration by attaching large-tip-opening electrodes to the skin on the head. **b**, The top four traces show jaw, pectoral fin, and operculum movements, as well as heart rate. The bottom two traces indicate fictive respiration and fictive swimming, which were recorded using electrodes placed on the head and tail, respectively, and could be distinguished using low-pass and high-pass filters. The blue fictive respiration trace was overlaid with the green jaw movement trace for comparison of respiration-coupled fictive electrical events, showing these electrical events can represent fictive respiration. Lines above the traces indicate swims and dots indicate respiration events; closed circles, respiration events during swims; open circles, respiration events outside swims. The boxed regions are shown at higher temporal resolution on the right. **c**, Left: Single trials (gray lines) and averages (colored lines) of craniofacial and fictive behaviors aligned by the onset of fictive respiration events that occurred not during (open circle) or during (closed circle) a swim bout. Right: Overlay of average craniofacial and fictive behaviors, aligned at the onset of fictive respiration events that were not during (open circles, top) or during (closed circles, bottom) swim bouts. The electrophysiological signals (blue) coincide with the physical movements associated with respiration, including movements of the jaw, fin, and operculum, confirming the electrical signals can be interpreted as fictive respiration. **d-f**, Respiration frequency (d), swim frequency (e) and swim vigor per bout (f) under normoxia and hypoxia in response to open-loop visual stimulation (16 fish), closed-loop visual stimulation (6 fish), or no visual stimulation (constant illumination of static dark/red stripes) (7 fish). Wilcoxon signed-rank test. **g**, Temporal traces showing respiration frequency, swim frequency, vigor per swim bout and individual swim bout duration during normoxia and hypoxia. The data was averaged in 30 second bins, and is shown as mean ± s.e.m. **h**, Histograms showing the distribution of inter-bout interval (left), vigor per bout (middle) and individual swim bout duration (right) for 29 fish. **i**, Spontaneous fluctuation of brain pO_2_ and fictive swimming in a paralyzed and a non-paralyzed fish. **j**, Left schematic indicates the brain pO_2_ metrics quantified in the bar plots on the right. Brain pO_2_ decreases due to a swim bout, then increases after swimming stops as O_2_ is replenished. Right, the minima of swim-induced brain pO_2_, the decay time from the pre-swim pO_2_ to 50% of ΔpO_2_ minima, the rise time from the pO_2_ minima to 50% ΔpO_2_ rise, and the time from swim stop to pO_2_ minima for 12 paralyzed and 4 non-paralyzed fish. Wilcoxon rank-sum test. **k**, Mean ± s.e.m delay between swim onset and the onset of resulting pO_2_ drops (indicated by the orange period in panel j) during normoxia and hypoxia from 6 paralyzed fish. **l**, Probability density plots of the durations of active swimming periods in paralyzed fish. Normoxia, black; hypoxia, blue. Mean durations are indicated as arrows on the top. **m**, Swim-triggered pO_2_ dynamics in paralyzed fish (centered at the onset of active swimming periods) during normoxia (black) and hypoxia (blue); mean ± s.e.m. are shown. Arrows indicate the mean durations of active swimming periods. During normoxia, pO_2_ fell below pre-swim levels at ~4 seconds (indicated as the gray period in panel j), similar to the mean duration (4.65 seconds) of active swimming periods. However, during hypoxia, the mean duration of swimming periods was only 2.31 seconds, whereas pO_2_ did not fall below pre-swim levels until over 8 seconds after swim onset, suggesting that a reactive model with a fixed pO_2_ threshold is not sufficient to describe behavior, and instead suggest a predictive neural control system (Fig. 1j). **n**, Schematic of the predictive model derived from an optimal control framework (Methods). In this formulation, brain pO_2_ dynamics follow a delayed linear dynamical system in which swimming induces a lagged decrease in pO_2_. Optimizing the system to avoid hypoxia while sustaining locomotion requires a series of computations. First, the model estimates future hypoxic stress by a multiplication of brain hypoxia and swim vigor, which can be matched to the oxygen-dependent swim encoding in NE-MO spiking (Fig. 3c,d). Next, the model sums this multiplication with brain hypoxia, which can be matched to oxygen-dependent tonic firing of NE-MO neurons (Fig. 3c, inner plot). Then, the model bases the control signal of behaviors on the temporal integration of the summed signal, which can be matched to astroglial dynamics. Finally, this control signal inhibits swimming. **o**, Example simulations of the model in (n) under normoxia (left) and hypoxia (right). Swim actions (1, swim; 0, no swim; black ticks) were matched to ‘swim’ in (n); pO_2_ profiles were matched to ‘brain pO_2_’ in (n); control signals were matched to ‘control signal’ in (n). Simulation shows an increase of the control signal, which lessens swim events during hypoxia.

**Extended Data Figure 2.**
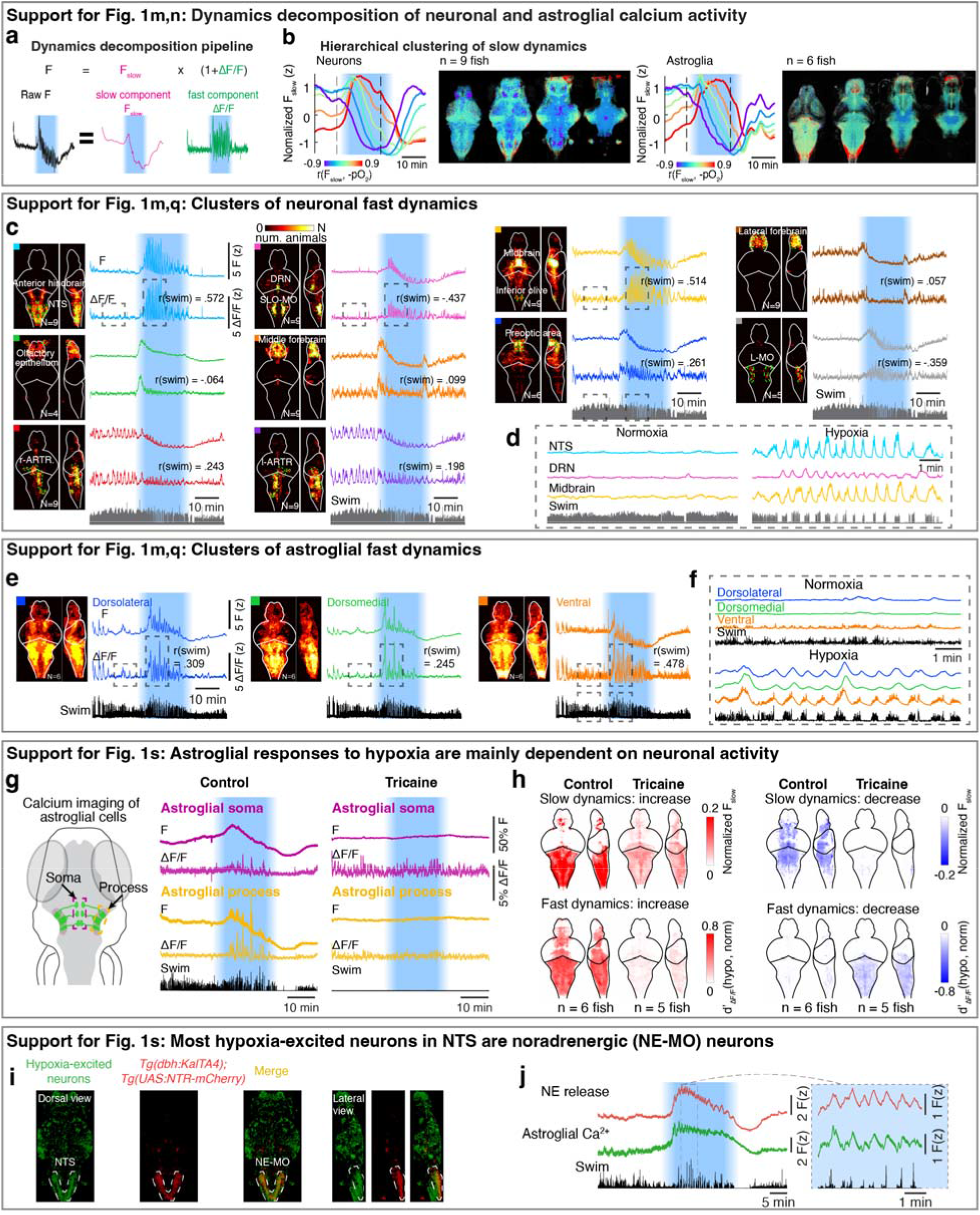
Hypoxia causes distinct changes to brain dynamics. **a**, Imaging data analysis pipeline showing decomposition of raw calcium activity into slow (F_slow_) and fast (ΔF/F) components, which were processed via rank correlation and functional clustering, respectively (Methods). **b**, Slow calcium components of neurons (left) and astroglia (right) show variable dynamics across different populations. We performed hierarchical clustering on slow components into 6 clusters and labeled the clusters by their rank correlation with brain pO_2_ change. Left: Temporal activity for each cluster; Right: spatial maps of 4 optical sections showing neurons that comprise each color-coded cluster. **c**,**d**, Left: Whole-brain maps for 10 clusters of neuronal fast calcium components across fish. N, number of fish that share the same cluster (9 fish in total; some clusters were missed in some fish due to imaging conditions). Right: calcium activity from the fish shown in Fig. 1m; top, z-scored raw fluorescence; bottom, z-scored fast components (ΔF/F). The colored square at the top left corner of each image indicates the brain region shown in the composite brain maps and correspond to the clusters with the same colors in Fig. 1q. r(swim) measures rank correlation between the instantaneous swim vigor and ΔF/F. Boxed regions of four neuronal clusters are shown at higher temporal resolution in (d). The NTS cluster and midbrain cluster are strongly correlated with swim vigor. The DRN cluster is strongly anti-correlated with swim vigor. **e**,**f**, Similar to (**c**) for astroglia. Whole-brain maps for 3 clusters of astroglial fast components across 6 fish. Right: calcium activity from the fish shown in Fig. 1m. Boxed regions are shown at higher temporal resolution in (f). **g**, Left: Schematic diagram showing GCaMP6f expression in astroglia (green), and regions of interest containing somata (purple box) and processes (yellow box). Right: Raw fluorescence (F) and fast calcium component (ΔF/F) in astroglial somata (purple) and processes (yellow) without or with tricaine (also known as MS-222, a sodium channel blocker) treatment. **h**, Whole-brain maps of astroglia slow and fast calcium dynamics in response to hypoxia without or with tricaine treatment. **i**, Colocalization of NTS neurons that are activated by hypoxia with NE-MO is shown using a *Tg(elavl3:H2B-jGCaMP7f);Tg(dbh:KalTA4);Tg(UAS:NTR-mCherry)* fish, in which all neurons express H2B-jGCaMP7f and noradrenergic neurons express mCherry. Green: hypoxia-excited neurons based on rank correlation between neuronal slow calcium component and brain pO_2_ change. Red: the expression of mCherry in noradrenergic neurons. Dashed line: the location of NE-MO cells in NTS. **j**, Hypoxia induces simultaneous slow and fast increases of NE release (red) and astroglial calcium (green) in the hindbrain of *Tg(elavl3:GRAB-NE2h);Tg(gfap:jRGECO1b)* fish. The dashed box region (left) is shown at higher temporal resolution (right).

**Extended Data Figure 3.**
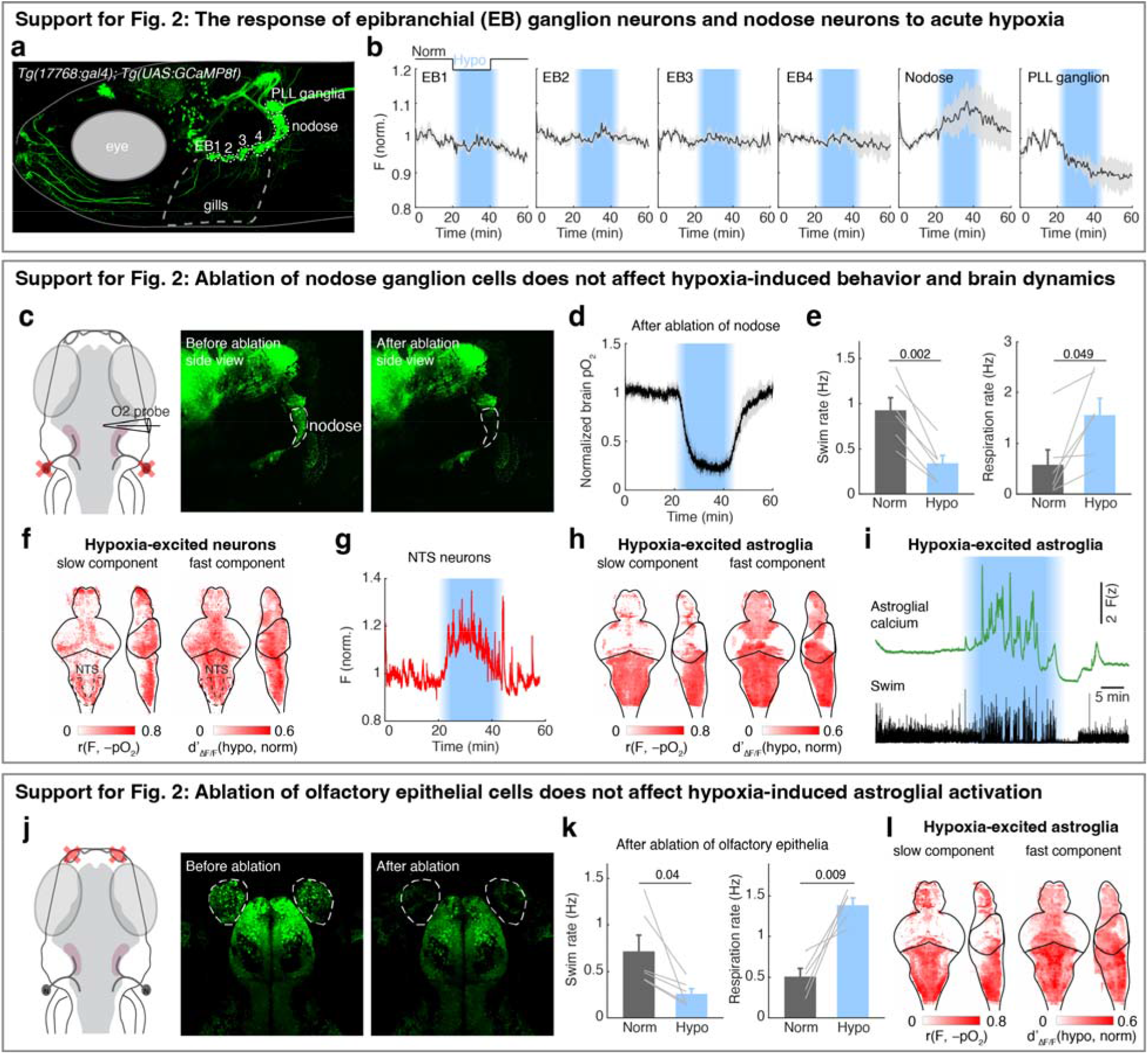
Peripheral chemosensory pathways that are not necessary for hypoxia-induced behavior and brain state changes. **a**, Fluorescent image of a 7-dpf *Tg(17768:gal4);Tg(UAS:jGCaMP8f)* zebrafish, in which epibrachial (EB) ganglia 1-4 (ganglia EB1/2 are also known as glossopharyngeal ganglia), nodose ganglia, and posterior lateral line (PLL) ganglia express jGCaMP8f. **b**, Normalized raw jGCaMP8f fluorescence of each cell population from (a) in response to hypoxia. Mean ± s.e.m. for 5 fish are shown. **c**, Schematic and fluorescent images showing the bilateral ablation of nodose ganglia in a *Tg(elavl3:H2B-jGCaMP7f)* fish. **d**,**e**, The change of brain pO_2_ (d), and swimming and respiration rate (e), in response to hypoxia in 6 nodose-ablated fish. Mean ± s.e.m for 6 fish are shown; Student’s paired t-test. **f**, Whole-brain dynamics maps showing the increase of slow and fast neuronal calcium components in response to hypoxia from 4 nodose-ablated *Tg(elavl3:H2B-jGCaMP7f)* fish. **g**, The average fluorescence of hypoxia-excited cells (red) in the NTS from a nodose-ablated *Tg(elavl3:H2B-jGCaMP7f)* fish. **h**, Whole-brain dynamics maps showing the increase of slow and fast astroglial calcium components in response to hypoxia from 6 nodose-ablated *Tg(gfap:GCaMP6f)* fish. **i**, Astroglial fluorescence in the hindbrain and swim events from a nodose-ablated *Tg(gfap:GCaMP6f)* fish in response to hypoxia. **j**, Schematic and fluorescent images showing the bilateral ablation of olfactory epithelial cells in a *Tg(elavl3:H2B-jGCaMP7f)* fish. **k**, Swim and respiration rate in response to hypoxia in 6 olfactory epithelia-ablated fish. Mean ± s.e.m for 6 fish are shown; Student’s paired t-test. **l**, Whole-brain dynamics maps showing the increase of slow and fast neuronal calcium components from 4 olfactory epithelium-ablated *Tg(gfap:GCaMP6f)* fish in response to hypoxia.

**Extended Data Figure 4.**
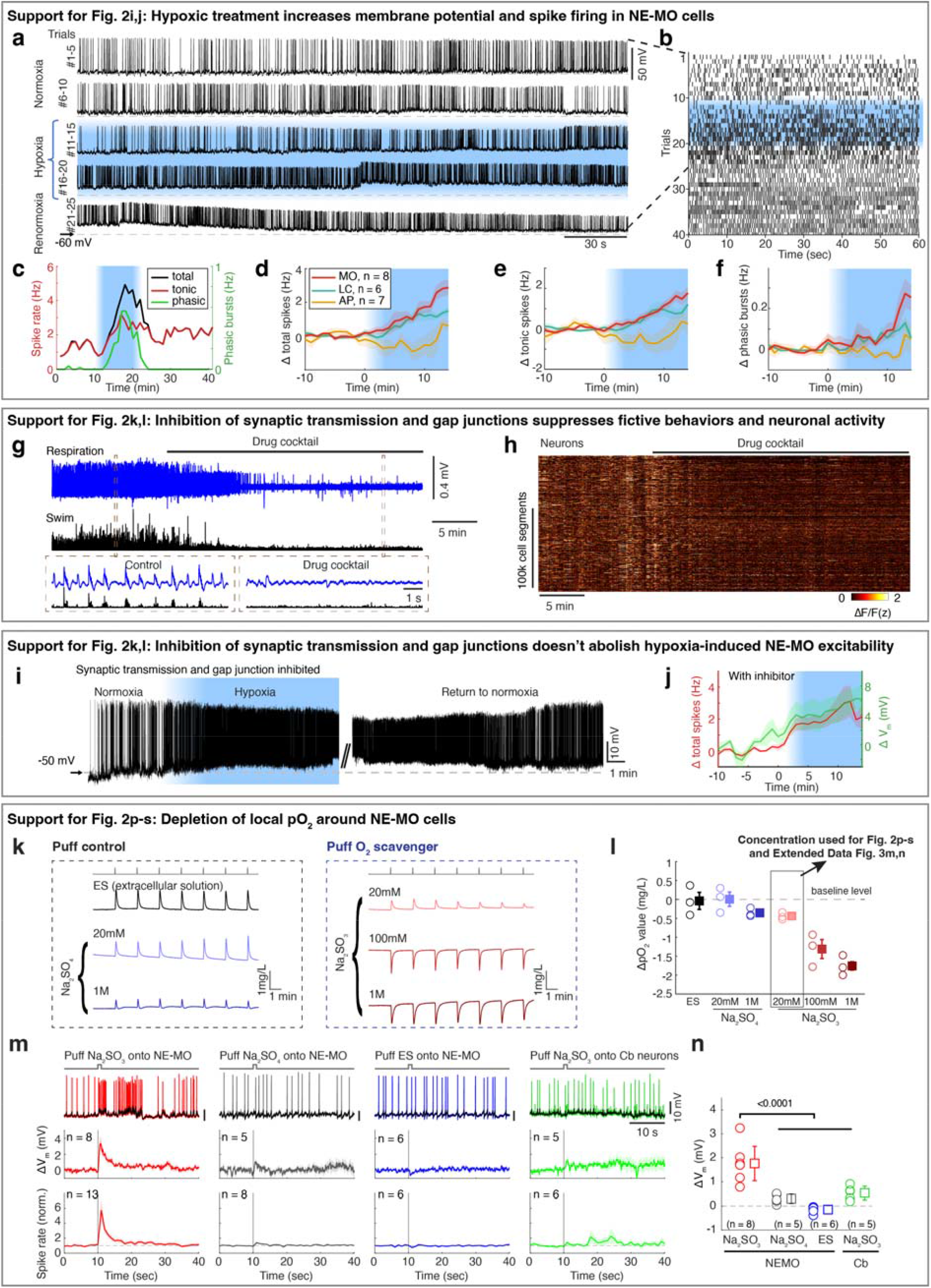
NE-MO is responsive to pO_2_ change. **a**,**b**, Example of raw electrical activity (a) and raster spikes (b) for the NE-MO cell shown in Fig. 2i, showing the change of membrane potential and spike firing during normoxia, hypoxia and return to normoxia. **c**, Time-series change of total, tonic spikes and phasic events of the NE-MO cell shown in (a,b). **d-f**, Mean ± s.e.m. change of NE-MO cell total (d), tonic (e) spike firing and phasic (f) events. The spike frequency change was calculated by subtracting the frequency during hypoxia from that during normoxia. n indicates the number of cells recorded. **g**, Fictive respiration (blue) and fictive swimming (black) during a 40-min recording (top) before and after treatment with a cocktail of drugs to block synaptic transmission and gap junctions. The boxed regions are shown at higher temporal resolution at the bottom. **h**, Neuronal jGCaMP7f fluorescence matrix from the fish in (g) showing the effect of the drug cocktail on spontaneous fast calcium (ΔF/F) dynamics across the whole brain. **i**, Example of NE-MO cell raw electrical activity showing the change of membrane potential and spiking under normoxia, hypoxia and return to normoxia. The experiment was performed with synaptic transmission and gap junction inhibited. **j**, Mean ± s.e.m. of total spiking (red) and membrane potential (green) for 7 NE-MO cells with synaptic transmission and gap junctions inhibited. n indicates the number of cells recorded. **k**,**l**, Example traces (k) and summary from 3 tests (l) showing the responses of an O_2_ sensor (Clark-type oxygen electrode) to puffing extracellular solution (ES), Na_2_SO_3_, or Na_2_SO_4_ onto the tip of the O_2_ sensor. Circles represent the average of 7 puffing-induced responses from 3 tests. Boxes are mean ± s.e.m.. The control solutions, ES and 20 mM Na_2_SO_4_, caused an upward deflection of apparent O_2_ level. This is likely due to a high shear force onto the O_2_ sensor caused by the high partial water pressure generated by puffs from the small diameter pipette tip (personal communication with specialists at manufacturer, Unisense). We therefore set the pO_2_ value caused by puffing ES as baseline. Thus, although puffing 20 mM of the O_2_ scavenger Na_2_SO_3_ caused a small upward response, the net pO_2_ change is about –0.4 mg/L after subtracting it by the baseline of puffing ES. In all O_2_ scavenger experiments, we used 20 mM Na_2_SO_3_ to get reliable responses across trials. **m**, Examples (top) showing spikes (colored traces) and V_m_ (black traces) in response to puffing Na_2_SO_3_, Na_2_SO_4_ or ES onto NE-MO, or puffing Na_2_SO_3_ onto cerebellar (Cb) neurons. Mean ± s.e.m. V_m_ (middle) and spikes (bottom) in response to each treatment for the indicated number of neurons. The vertical lines at t=10 seconds indicate puffs. Na_2_SO_4_ only caused a small increase in V_m_ and spike rate. **n**, Quantification of puff-induced V_m_ change, based on data from the middle row in (**m**). Circles represent individual neurons. Boxes represent mean ± s.e.m. The numbers in (**m**,**n**) indicate the number of neurons quantified; Wilcoxon rank-sum test.

**Extended Data Figure 5.**
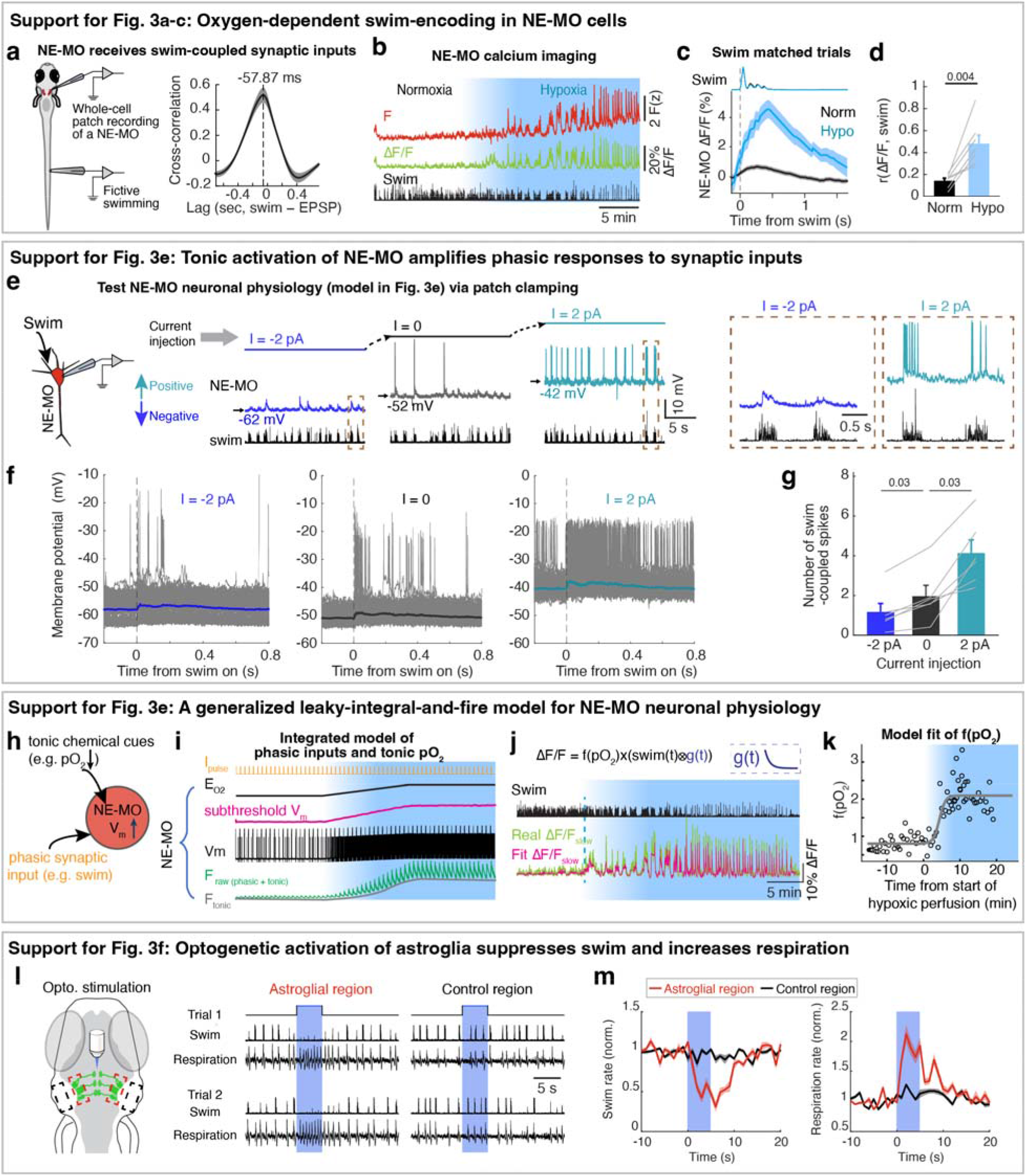
Responsiveness of NE-MO to synaptic inputs is gated by its resting membrane potential. **a**, Left, schematic of simultaneous whole-cell patch-clamp recording of a NE-MO cell and electrophysiological recording of fictive swimming. Right, cross-correlation between swim signals and EPSPs of the NE-MO cell in Fig. 3a; the −58 ms lag from EPSP to swim signal indicates that synaptic input to the NE-MO neuron occurs 58□ms after the swim bout. Mean ± s.e.m. across trials is shown; a dashed line indicates the timing of the peak of the cross-correlation. **b**, Example traces showing raw calcium fluorescence (F, top) and fast calcium component (ΔF/F) of NE-MO neurons, as well as fictive swimming (bottom), in a *Tg(th1:gal4);Tg(UAS:GCaMP6f)* fish. **c**, Averaged swim-triggered fast calcium activity (ΔF/F) of NE-MO cells in (b) when similar swim bouts were selected under normoxia (black trace) and hypoxia (blue trace). Top, swim traces in the matched trials. Mean ± s.e.m. are shown. **d**, Mean ± s.e.m. of Spearman correlation coefficient between swim vigor and NE-MO fast dynamics under normoxia and hypoxia. Wilcoxon signed-rank test across 7 fish. **e**, Schematic and example traces showing NE-MO activity and swim events when NE-MO was held at different V_m_ by whole-cell patch-clamp recording. Positive or negative current injection was applied to mimic the change of oxygen-dependent parameter V_m_. **f**, Overlay of individual swim-coupled trials (gray) and average of all trials (colored line) in each current injection condition. **g**, Bar plots of swim-coupled spikes at different current injection conditions. The number of swim-coupled spikes increases with V_m_. Mean ± s.e.m. for 6 cells are shown. Wilcoxon signed-rank test. **h-i**, Biophysical cellular model showing the integration of oxygen levels and motor signals at NE-MO; (h) model schematic; (i) simulation. We assumed that oxygen impacts NE-MO cell physiology via a parameter E_O2_ (which resembles the reversal potential in a generalized leaky integral-and-fire model; Methods). E_O2_ increases with hypoxia levels, resulting in a change of the subthreshold voltage, V_m_, which mimics real data. To capture the change of phasic spikes of the cell, we simulated the synaptic inputs as pulse currents, I_pulse_. The spiking output of the cell is shown as the dynamics of V_m_ and that for GCaMP fluorescence is shown as F_raw_, which can be decomposed into F_slow_ and ΔF/F (that responds to I_pulse_). **j**, A time-series fit of NE-MO ΔF/F through the transform parameter f(pO_2_) and instantaneous swim vigor during normoxia and hypoxia. We examined the model in calcium dynamics where we assumed that swim vigor was the synaptic input, and NE-MO ΔF/F was the output. We thus fitted a linear model (see Supplementary Methods) of the swim vigor (that was filtered through an exponential decay function of g(t)), indicating the decay of NE-MO GCaMP6f fluorescence to ΔF/F every 30 seconds. The oxygen-dependent gain factor f(pO_2_) was measured by the linear coefficient of the fit. Green and magenta traces indicate real ΔF/F data and fitted fast activity based on swim vigors (variance explained, 79.63%), respectively. **k**, Dynamics of oxygen-dependent gain factor f(pO_2_) during the normoxia-hypoxia transition, which agrees with the oxygen-dependent swimming encoding in Fig. 3c,d. **l**,**m**, Examples of an individual fish (l) and mean ± s.e.m. for 5 fish (m) showing that pulsed (10-ms pulses at 10-Hz for 5 seconds, blue shading) optogenetic activation of astroglia (dashed red boxes, left in **l**), but not a control region (dashed black boxes, left in l), using *Tg(gfap:CoChR)* fish induces behavioral phenotypes that are similar to those induced by hypoxia and to stimulation of NE-MO neurons (Fig. 1e).

**Extended Data Figure 6.**
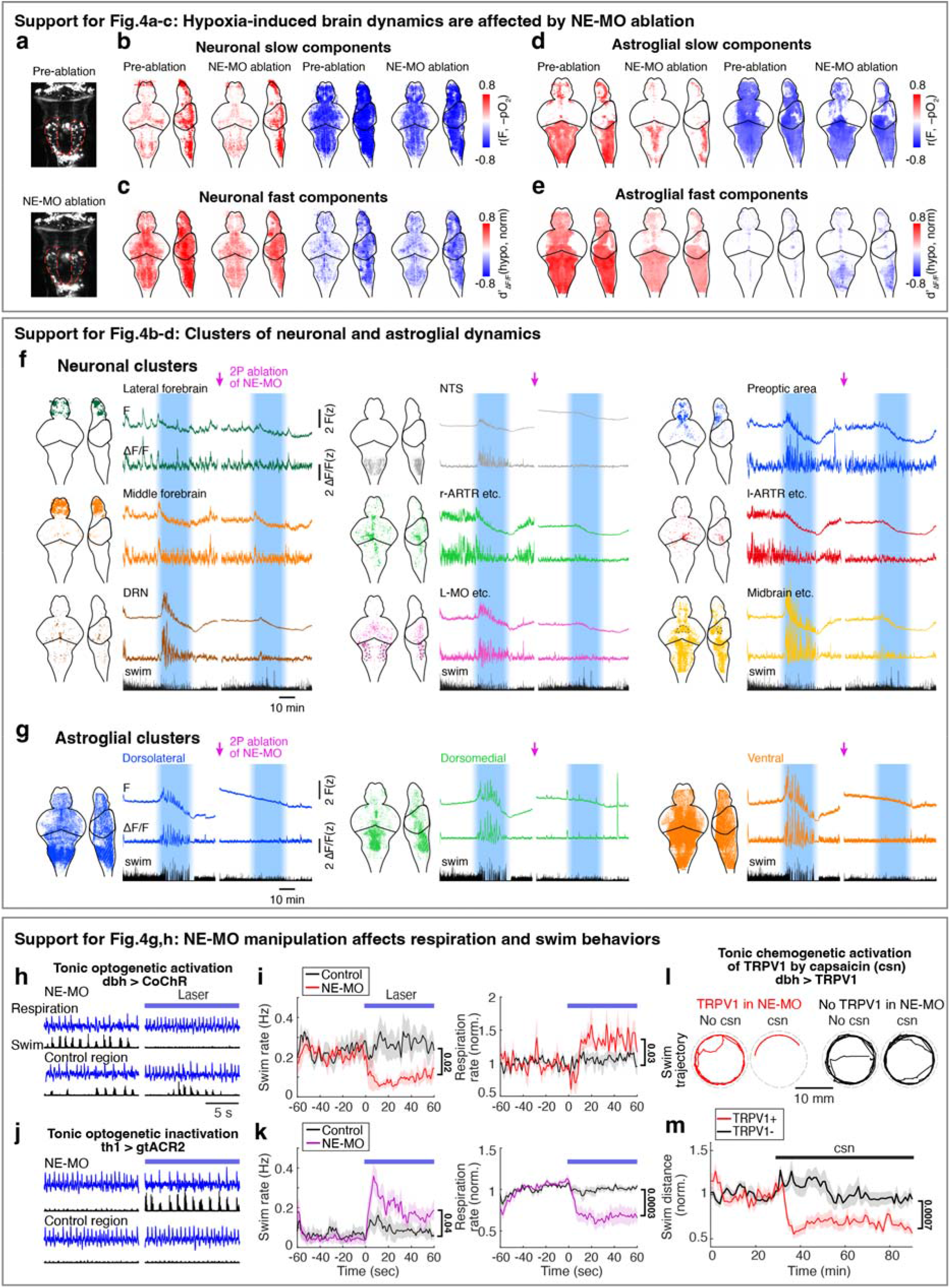
Noradrenergic signaling is important for hypoxia-induced brain dynamics and behavioral phenotypes. **a**, Fluorescent images of *Tg(th1:Gal4);Tg(UAS:NTR-mCherry)* fish before and after two-photon ablation of NE-MO while performing whole-brain calcium imaging under the same light-sheet microscope. The dashed red line indicates ablated regions. **b-e**, Whole-brain maps for hypoxia-induced slow and fast calcium dynamics before and after NE-MO ablation for neurons (5 fish, b,c) or for glia (8 fish, d,e). Red and blue colors indicate hypoxia-excited and hypoxia-inhibited cells, respectively. Hypoxia-induced increase of slow and fast astroglial dynamics is largely reduced after NE-MO ablation, which indicates a brain-wide modulatory role for NE. **f**,**g**, The brain map, average fluorescence (F), and fast component activity (ΔF/F) of neuronal (f) and astroglial (g) clusters from the example fish in Fig. 4b,c, showing that NE-MO ablation reduced the calcium responses to hypoxia in many astroglial and neuronal populations. **h**,**i**, Examples of individual fish (h) and mean ± s.e.m. for 5 fish (i) showing that continuous (60 seconds, blue line) optogenetic stimulation of NE-MO using *Tg(dbh:KalTA4); Tg(UAS:CoChR-eGFP)* fish induces effects on swimming and respiration that are similar to those observed in hypoxia. The control region of laser illumination was located outside of the brain. The stimulation areas for NE-MO and control are similar. Student’s paired t-test. Due to variable swim and respiration rates in some of these fish, some graphs show normalized data rather than actual rates. **j**,**k**, Examples of individual fish (j) and mean ± s.e.m. for 7 fish (k) showing that continuous (60 seconds, blue line) optogenetic inhibition of NE-MO using hypoxic *Tg(th1:gal4); Tg(UAS:GtACR2-eYFP)* fish induces phenotypes that are opposite to those induced by hypoxia. Student’s paired t-test. **l**,**m**, Example 5-minute swim trajectories for single fish (l) and mean swim distance for 45 TRPV1-positive and 45 TRPV1-negative fish (m) showing that chemogenetic stimulation of noradrenergic neurons, including the NE-MO cluster, using capsaicin treatment of *Tg(dbh:KalTA4);Tg(UAS:TRPV1:mCherry)* fish induced hypoxia-like suppression of swimming. Student’s unpaired t-test.

**Extended Data Figure 7.**
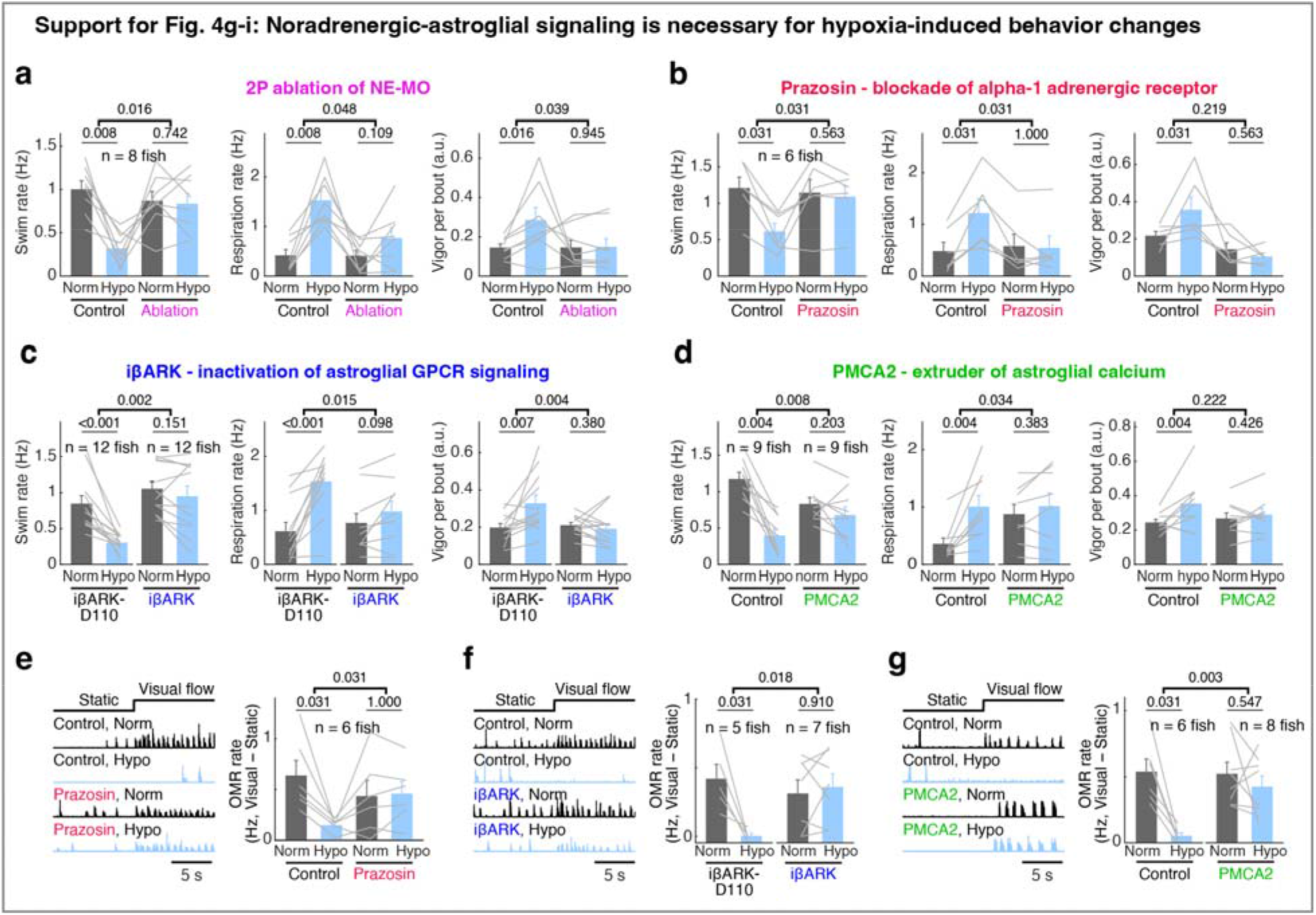
Noradrenergic-astroglial signaling is necessary for hypoxia-induced coping behaviors. **a-d**, Effects of NE-MO 2P ablation (a), prazosin treatment (b), or expression of iβARK (c) or hPMCA2 (d) in astroglia on hypoxia-induced changes in swim rate, respiration rate and swim vigor. The control group in (c) used *Tg(gfap:iβARK-D110)* fish, which expresses a mutant transgene that does not inhibit Gq GPCR signaling. **e-g**, Effects of prazosin treatment (e), or expression of iβARK (f) or hPMCA2 (g) in astroglia, on hypoxia-induced optomotor responses, which was calculated by subtracting swim rate during visual stimulus to that during static period. Gray lines in (a-g) indicate individual fish and bar graphs indicate mean ± s.e.m. Wilcoxon signed-rank test for paired statistics in same groups, Wilcoxon rank-sum test for unpaired statistics between groups.

**Extended Data Figure 8.**
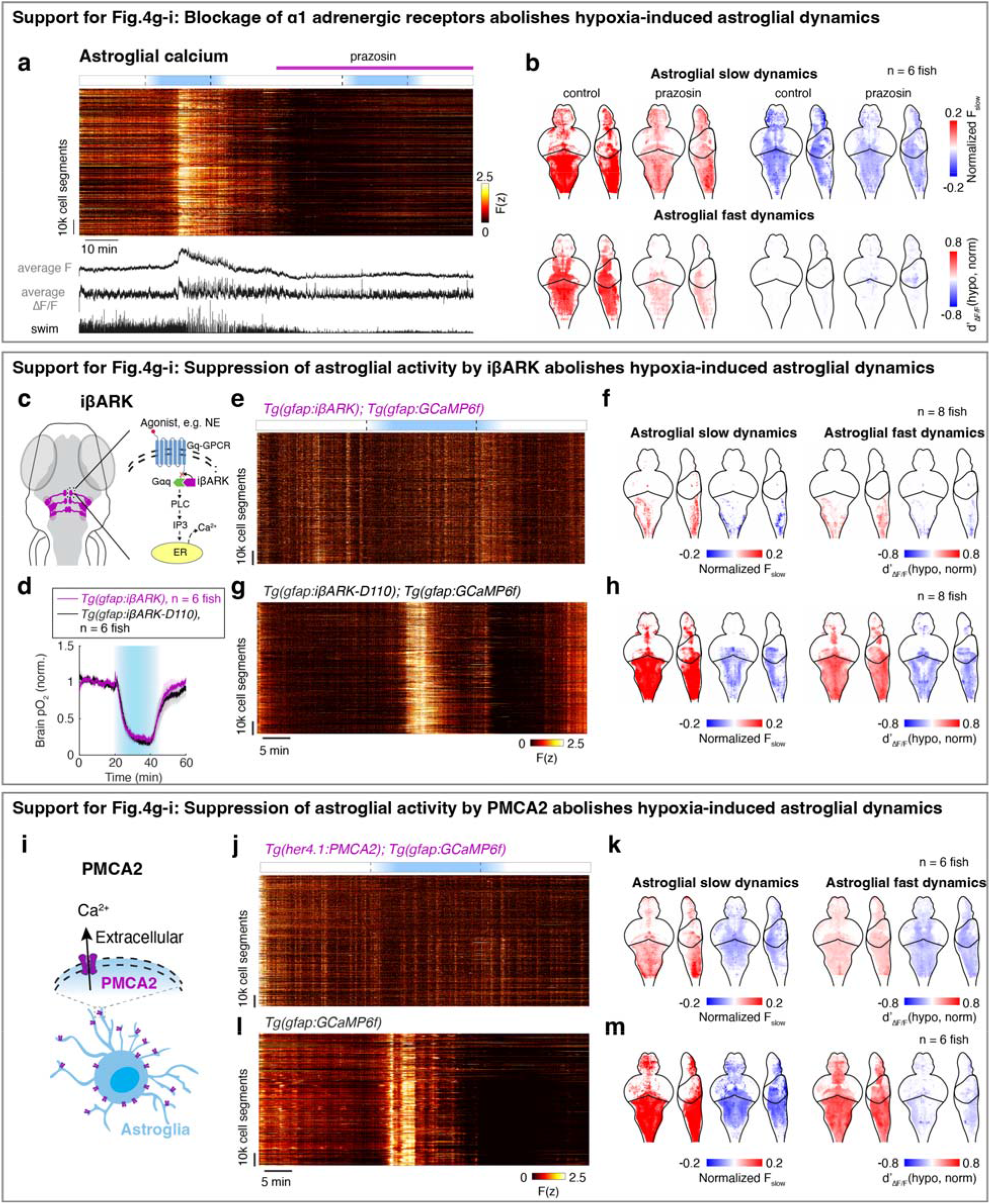
Inhibition of astroglial activity suppresses the effects of hypoxia on astroglial dynamics. **a**, Whole-brain astroglial raw fluorescence matrix (top) and average calcium activity and fictive swimming (bottom) showing the effect of prazosin treatment on hypoxia-induced astroglial activity and swim behavior. **b**, Whole-brain maps for slow and fast calcium activity in astroglia before and after prazosin treatment for 6 fish. **c**, Schematic showing the Gq GPCR signaling pathway in astroglia. Expression of an artificial peptide (iβARK) blocks the activation of Gq by GPCR agonists, e.g., NE. **d**, Brain pO_2_ in response to hypoxia in *Tg(gfap:iβARK)* (purple) and *Tg(gfap:iβARK-D110)* (black) fish. Mean ± s.e.m. for 6 fish are shown. **e-h**, Fluorescence matrix (e,g) and whole-brain maps (f,h) showing effects of hypoxia on astroglial calcium activity in *Tg(gfap:GCaMP6f);Tg(gfap:iβARK)* (e,f) and *Tg(gfap:GCaMP6f); Tg(gfap:iβARK-D110)* (g,h) fish. Expression of iβARK suppresses astroglial fast and slow calcium dynamics in response to hypoxia but expression of the negative control protein *iβARK-D110* does not. **i**, Schematic showing that PMCA2 functions as a calcium extruder that pumps calcium out of the cytoplasm when expressed in astroglia. **j-m**, Fluorescence matrix (j,l) and whole-brain maps (k,m) showing effects of hypoxia on astroglial calcium activity in *Tg(gfap:GCaMP6f);Tg(her4*.*1:PMCA2)* (j,k) and *Tg(gfap:GCaMP6f)* (l,m) fish. Expression of *PMCA2* suppresses astroglial fast and slow calcium dynamics in response to hypoxia.

## Supplementary information

### Supplementary Methods

#### Cellular model for NE-MO dynamics

We used a generalized leaky integrate-and-fire model, GLIF (Teeter et al, 2018), to simulate the NE-MO cell in the whole-cell patch-clamp recordings (Fig. 3e; Extended Data Fig. 5h,i). The model was made up of two parts: a linear dynamics of voltage change to input and a dynamics of the firing threshold:

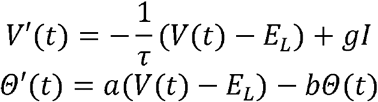

*E*_*L*_ is the resting membrane potential, *τ* = 20 ms, *b* = 0.009 /ms are time constants of membrane potential and voltage dependence of the threshold, respectively, and *α* - 0.0001 /ms couples the membrane potential to the threshold. *I* is the external current and *g*=1 μS is the impedance.

If *V*(*t*) > *Θ*(*t*) + *Θ* _∞_, a spike is generated, and then *V*(*t*) was reset as follow:

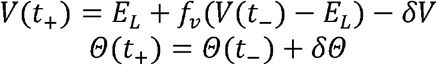

where *f* _*ν*_ = 0.6, and *δV* = −5 mV, are the slope and intercept of the linear relationship of the voltage before and after a spike, respectively; *Θ*_∞_ = −39 mV is the baseline firing threshold; the transient increase of the firing threshold after a spike event *δΘ* = 4 mV reflects the repolarization or after-hyperpolarization which inhibits spike generation after a spike.

In the simulation, we used *E*_*L*_ = −61 mV for normoxia and *E*_*L*_ = −57 mV for hypoxia, respectively. A linear increase of *E*_*L*_ from −61 mV to −57 mV presents the dynamics of oxygen level transiting from normoxia to hypoxia. Tonic spikes were measured when external input *I* - 0 pA, while phasic spikes were measured when a transient pulse *I* - 9 pA was provided for 300 ms every 3 seconds.

#### Cellular model fits to fluorescence dynamics

We fitted our cellular model to fluorescence dynamics (Extended Data Fig. 5j,k), Δ*F*/*F*(*t*), every 1 minute:

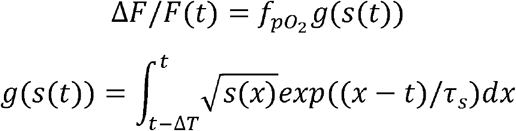

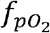 is the transform scalar (which depends on O_2_ level) from swim to fluorescence; *g*(·) is an exponential filter function of the square root of swim power in a time window Δ*T* before the fluorescence. The performance of the fitting was evaluated using variance explained as

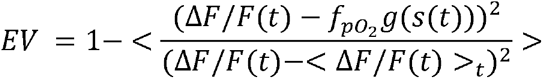

where <·> is the arithmetic average.

#### Brain-body control system model

We include an alternative model of how external oxygen (O_2_) replenishes blood O_2_, which in turn supplies muscle O_2_ through circulation which leads to qualitatively similar solutions as presented in the main text. Let X, M, D denote blood O_2_, muscle O_2_, and instantaneous oxygen demand. Their stochastic dynamics are:

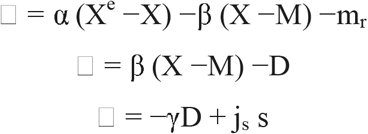

where sdt ~ Poiss(u) represents the probability of swimming, given a control rate u. Here, α and β describe oxygen exchange between environment–blood and blood–muscle, γ is the decay of oxygen demand, m□ is metabolic rate, j□ the swim intensity, and X^e^ the external O_2_ level.

The system aims to keep blood O_2_ near an optimal setpoint X* while minimizing deviations from a preferred swimming rate u*:

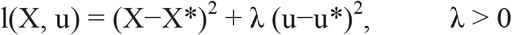

Approximating s by its mean rate u and defining centered variables X□ = X−X*, û = u−u*, the system becomes affine-linear. The optimal controller is of LQR form (Anderson & Moore, 2007):

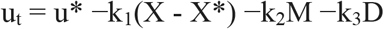

The constants k_1_, k_2_, k_3_ depend on the parameters α, β, γ. In our model simulations, we found k_3_ ≈ 0 consistently, and we will therefore exclude it and simplify the controller as

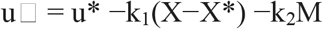

The controller directly senses blood O_2_ (X) and demand (D) but not muscle O_2_ (M), making the system only partially observable. For partially observable systems, the optimal strategy obeys the separation principle, where the optimal controller consists of an optimal state estimation, such as the Kalman filter, and an optimal deterministic control. Therefore, we postulate that the missing variable M can be approximated by a leaky integration of D:

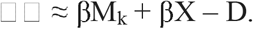

M_k_ is the estimated muscle oxygen. Finally, to prevent unbounded control, we introduce a sigmoid nonlinearity approximately linear near equilibrium:

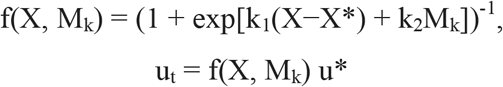

This formulation captures a simple closed-loop strategy for oxygen homeostasis under stochastic behavioral drive with signals X and M.

## Supplementary Videos

**Video S1: Animated description of Figure 3f**

